# Energy deposition and receptor saturation set a tumour size threshold for radiopharmaceutical therapy: insights from a pharmacokinetic–tumour dynamics model

**DOI:** 10.1101/2025.05.28.656722

**Authors:** Elahe Mollaheydar, Babak Saboury, Arman Rahmim, Eric N. Cytrynbaum

**Affiliations:** School of Biomedical Engineering, University of British Columbia, Vancouver, BC, Canada; Department of Integrative Oncology, BC Cancer Research Institute, Vancouver, BC, Canada; Institute of Nuclear Medicine, Bethesda, MD, US; Departments of Radiology and Physics, University of British Columbia, Vancouver, BC, Canada; Department of Mathematics, University of British Columbia, Vancouver, BC, Canada

**Keywords:** Radiopharmaceutical Therapy, Tumour Dynamics

## Abstract

Radiopharmaceutical therapies (RPTs) offer targeted radiation delivery to tumour cells, yet treatment outcomes vary substantially across patients while dosing protocols remain largely uniform. Translating mechanistic insight into improved protocols requires models that couple radioligand (RL) pharmacokinetics, spatial tumour biology, and radiation response — a combination that existing models have not yet fully achieved. We coupled a cellular automaton for spatially resolved tumour dynamics to a pharmacokinetic compartment model that tracks RL from injection through receptor binding and internalization. We estimate the energy deposited by radiation emitted from RL within the tumour and use a linear-quadratic radiobiological survival probability to determine the impact of the treatment. Simulating heterogeneous tumours across a range of conditions, we find that treatment outcome is governed primarily by tumour size and receptor expression levels and is relatively insensitive to the injected amount per cycle but highly dependent on inter-injection time intervals. The model’s uniform RL delivery separates radiobiological resistance from delivery effects — two mechanisms that are spatially correlated in real tumours — and the simulations suggest that compromised RL delivery may play a larger role in hypoxic treatment failure than radioresistance alone. These findings provide a mechanistic basis for patient stratification by receptor expression, yield a new size-based rationale for multi-injection protocols, and demonstrate that spatially resolved modelling can reveal treatment principles inaccessible to non-spatial approaches.

## Introduction

Radiopharmaceutical therapies (RPTs) are emerging as a very promising paradigm for treating various cancers by delivering targeted radiation doses to tumour cells while sparing healthy tissues [1]. RPTs rely on the unique properties of radiopharmaceuticals, consisting of a radionuclide combined with a targeting vector, such as a peptide or antibody. These vectors selectively bind to cancer-specific markers, enabling precise delivery of cytotoxic radiation [2]. Over the past few years, RPTs have gained significant attention in the treatment of metastatic cancers, where conventional treatments may be inadequate [1, 3]. They have demonstrated great potential, specifically for neuroendocrine tumours and prostate cancer, as shown by successes with Lutetium-177 (Lu-177) [4, 5, 6] and Actinium-225 (Ac-225) [7] labeled therapies.

In RPT practice, a ‘one-size-fits-all’ approach to dosage and timing of injections is commonly deployed, neglecting patient-specific differences in pharmacokinetics that can produce variable delivered doses [8, 9]. The growing body of evidence supporting the efficacy of RPTs [10, 11, 12] highlights the need for a deeper understanding of the key factors that influence treatment personalization — the process of tailoring treatment to the individual characteristics of the patient and their tumour. Patient-specific factors relevant to RPT outcomes include tumour size and vascularity, receptor expression levels, and individual pharmacokinetics governing how radioligand is distributed and cleared. On the treatment design side, the adjustable parameters include the total injected activity, the number of injections and the interval between them, and the choice of radioligand and its targeting vector. Advances in quantitative imaging and dosimetry have made it increasingly feasible to measure these patient-specific factors [13], paving the way for dosing protocols adapted to the individual. Theranostic platforms pair diagnostic imaging with therapy using the same molecular target (e.g., the PSMA protein in prostate cancer) [14, 15], offering a particularly direct route to treatment individualisation [16, 17].

Mathematical and computational models can illuminate the mechanistic basis of RPT outcomes and predict how treatment parameters — injected activity, injection timing, tumour biology — interact to determine whether treatment succeeds or fails. Among various modelling approaches, cellular automata (CA) have emerged as an effective tool for capturing spatial tumour heterogeneity at cellular resolution [18, 19, 20, 21]. A parallel tradition of compartmental PBPK models has examined how tumour-level quantities — total tumour volume, receptor expression, ligand affinity, and internalization rate — govern RL distribution and treatment outcome in ^177^Lu-PSMA-617 therapy [22, 23, 24]. At finer spatial scales, Birindelli et al. [25] coupled histology-derived microvascular networks with a convection– reaction–diffusion model of RL transport and LQ-based cell survival to show that hypoxic, poorly vascularized regions receive subtherapeutic radiation doses in ^177^Lu-PSMA-617 therapy, implicating RL delivery failure as a key resistance mechanism. More recently, spatially resolved models of radioligand transport in solid tumours have begun to appear [26], though coupling spatial RL delivery to tumour growth and radiation response in a unified framework remains largely unaddressed.

The radiobiological components of the present work draw on established modelling traditions: partial differential equation models of spatially distributed tumour cell populations under radiotherapy [27]; tumour control probability formulas derived from cell-cycle models distinguishing radiosensitive and radioresistant subpopulations [28]; and analyses of how repopulation kinetics within the LQ formalism inform fractionation schedule design [29]. Here, we address the gap between these traditions with a model that couples spatially resolved tumour dynamics and oxygenation with radioligand pharmacokinetics and LQ radiation response, establishing a theoretical basis for more efficient and personalized RPT protocols.

We developed a hybrid discrete-continuous stochastic-deterministic model to understand the effect of cancer treatment using the radiopharmaceutical ^177^Lu-PSMA-617 with varying injection protocols on heterogeneous tumours with varying receptor density. The full model includes a stochastic spatial CA to describe tumour growth and dynamics, a partial differential equation (PDE) for oxygen delivery, diffusion, and uptake at the cell level, a pharmacokinetic (PK) component consisting of a system of ordinary differential equations (ODEs) to track the amount of radioligand (RL) across several compartments over time, and a pharmacodynamic (PD) component describing the impact of RL on tumour cells.

The components are coupled in various ways, as illustrated in Fig. 1. Tumour dynamics involve processes that depend on local oxygen (cell division rate, state transitions between normoxic, hypoxic, and necrotic) and on the radiation exposure (induction of apoptosis). Radiation exposure is determined by the delivery (PK), decay, and absorption (PD) of RL. The tumour cell population can influence the delivery of oxygen and RL by occluding vessels and also influence the amount of RL present in the tumour by dictating the amount of RL receptors which is proportional to the tumour population size. We describe the four components of the model and their coupling in more detail in the Methods section.

**Figure 1.**
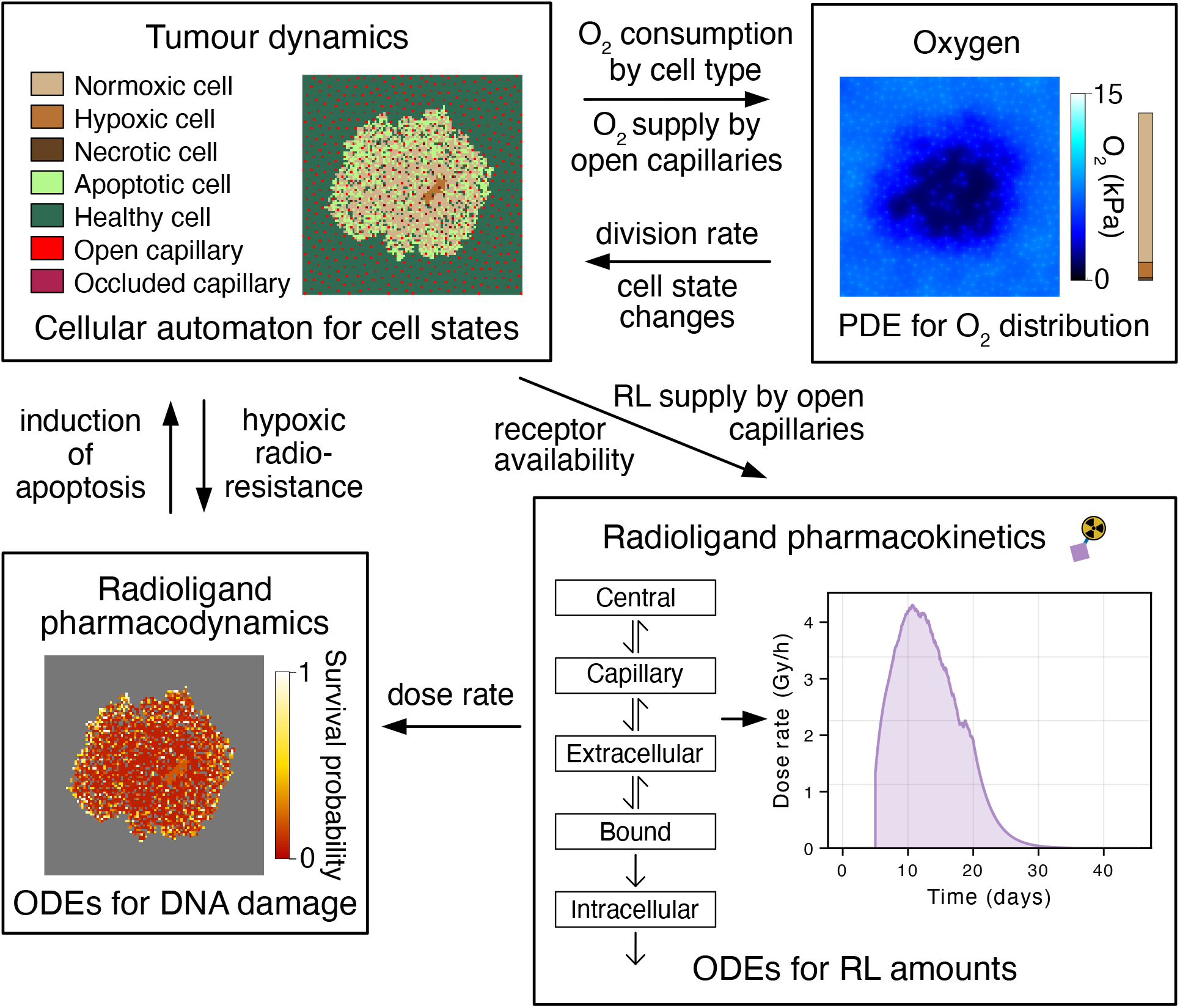
Model overview. Each box summarizes a submodel with titles corresponding to subsections of the Methods section. Arrows between boxes indicate the influence of one submodel on the other. For example, the arrow labeled “division rate” indicates that the oxygen concentration from the oxygen submodel influences the division rate of cells in the tumour dynamics submodel. The same arrow is also labeled “cell state changes”, indicating that transitions between normoxic, hypoxic, and necrotic are controlled by the oxygen concentration. The three tumour images are snapshots from the same moment (day 48) of a single simulation. The “Survival probability” in the pharmacodynamics box, indicating the colour scale, is the Lea-Catcheside survival fraction which is determined by the dose rate and cell type (hypoxic cells are radioresistant) and used as a stochastic survival probability in the CA. The text at the bottom of each box indicates the type of model used for that submodel (e.g. ODEs, PDEs, or CA).

A factor that requires attention in coupled treatment-response models, and plays an important role in this study, is the fraction of radiation energy emitted from the tumour that is absorbed within the tumour itself and not lost to the surrounding tissue. This quantity is a special case of the absorbed fraction in the MIRD (Medical Internal Radiation Dose) formalism where source and target are the same region, specifically, the tumour. We refer to it here as the intratumoral energy deposition fraction (EDF) to emphasize its role as a size-dependent variable governing treatment efficacy. It is described briefly in the Methods section and in more detail in the Appendix and Supplemental Information. Because the EDF decreases sharply with tumour size, small tumours receive a disproportionately low effective dose even when radioligand delivery is unimpaired. We incorporate this size-dependent EDF into the model and find that it produces a threshold tumour size below which treatment fails — a prediction that runs counter to the prevailing focus on treatment failure in large, hypoxic tumours. Our model’s uniform RL delivery, while a simplification that limits its applicability to larger tumours with compromised vasculature, has the advantage of isolating the radiobiological effects of hypoxia from delivery effects, two mechanisms that are difficult to separate experimentally.

## Results

### Tumour growth in the absence of treatment

We simulated a growing tumour without RL injection to illustrate and characterize the tumour dynamics thereby providing a baseline for tumour growth, capillary occlusion, and the development of hypoxia in the absence of treatment.

Four snapshots of a growing tumour are shown in Fig. 2. The initial tumour consisted of four normoxic cells. Over 80 days, it grew to about 1 mm in diameter (thousands of cells in the plane), during which time numerous capillaries (red) on the interior were occluded (dark red) leading to the appearance of a large hypoxic region and later a smaller necrotic core (see legend for colours). Capillaries were spread uniformly throughout the domain with a typical spacing of 3–5 cell lengths (605 capillaries per mm^2^). The (dark red) occluded capillaries align roughly with the hypoxic and necrotic regions.

**Figure 2.**
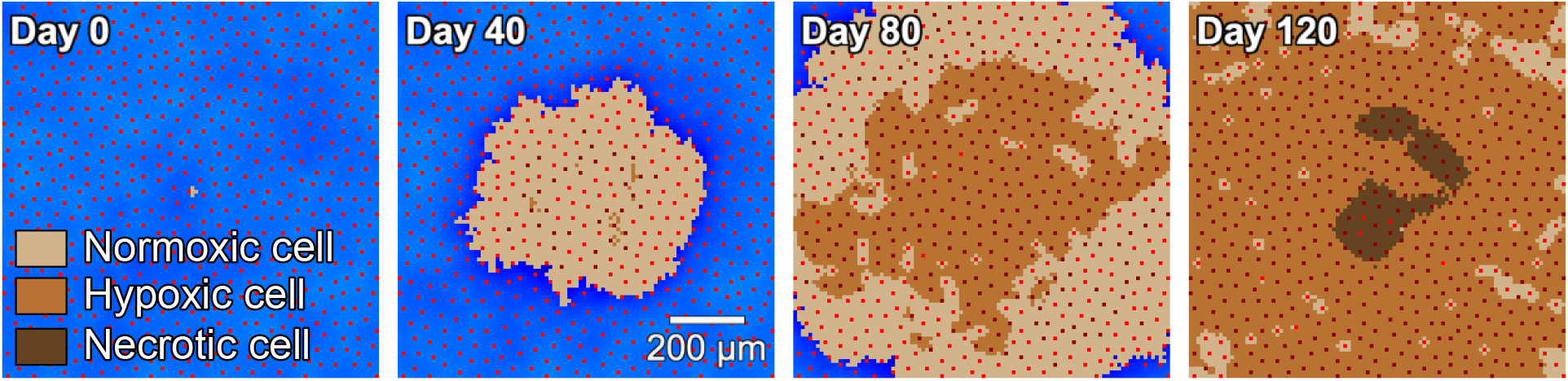
Tumour growth without treatment. Starting with a small and fully normoxic (light brown) tumour consisting of four cells, some capillaries get occluded and hypoxic regions (darker brown) begin to appear; later, necrotic patches (darkest brown) appear. The tumour is shown over the oxygen field (blue) with the oxygen partial pressure replacing the healthy cells (green) shown in the analogous example image in Fig. 1. These images are magnified by a factor of four for clarity (e.g. so that the state of capillaries are visible), showing a 1 mm *×* 1 mm region at the centre of the 4 mm *×* 4 mm domain. The full oxygen fields (not shown) look similar to the one shown in Fig. 1, with low oxygen at the edges of the tumour dropping even lower at the centre.

### Tumour size determines treatment outcome through the energy deposition fraction

Tumour size has been implicated in treatment success through various pathways, including compromised oxygen and RL delivery due to vessel occlusion and loss of beta particles from RL decay at the periphery of the tumour to surrounding healthy tissue which we quantify using the energy deposition fraction (EDF - see Methods for details). We carried out simulations under conditions with minimal capillary occlusion and hypoxia to focus on the impact of the EDF.

We simulated three tumours of different sizes undergoing treatment, illustrated in Fig. 3. Conditions, aside from initial size, were the same in all three simulations but responses to treatment differed. Treatment failed for the smallest tumour (initially four cells or roughly 28 *µ*m diameter) due to the survival of some cells that continued to grow after treatment. The intermediate- (initially 28 cells or roughly 70 *µ*m diameter) and large-sized tumours (initially 2648 cells or 600 *µ*m diameter) responded well, being eliminated by day 13 and 16 respectively. Although the injected amount was the same for all three, the EDF (*f*_deposit_) modulated the dose significantly — the vast majority of beta particles emitted from the small tumour were deposited outside the tumour. The factors for the three tumours were 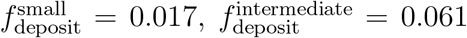, and 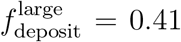, introducing a large difference in effective doses between the tumours.

**Figure 3.**
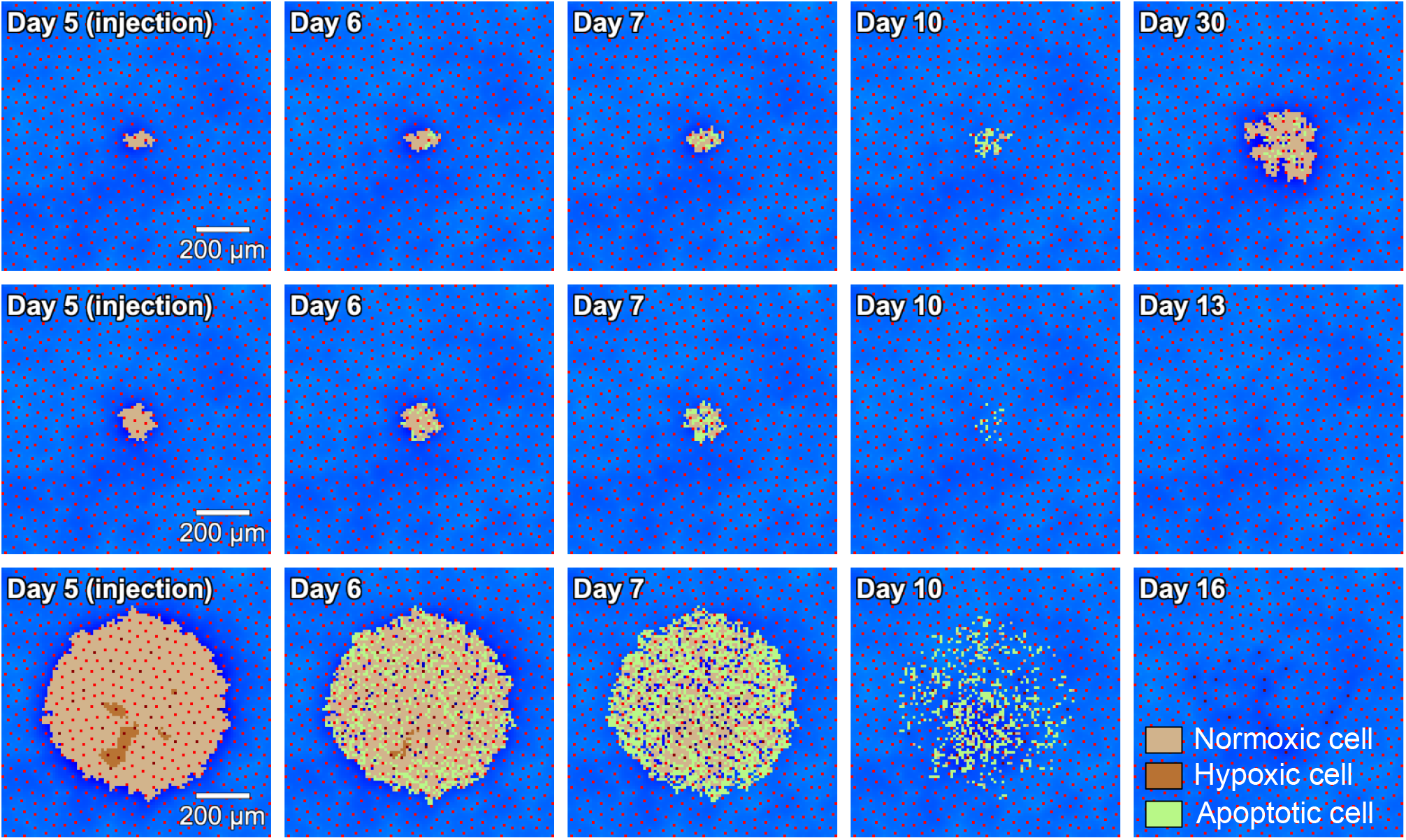
Three growing tumours of different sizes but otherwise the same conditions under RPT treatment with minimal hypoxia present (in the large tumour only). Top row: Treatment of a small tumour (initially four cells) failed. Middle row: The intermediate-sized tumour (initially 28 cells) was just above the size threshold of treatment success upon injection for the given conditions. Bottom row: For the largest tumour tested (600 *µ*m diameter, 2648 cells), treatment was successful and displayed a similar time-course to the intermediate-sized tumour. For the intermediate and large sized tumours, the final panel shows the first day on which no tumour cells remained. As described in the text, these differing responses can be explained by the energy deposition fraction (EDF).

The two smaller tumours were too small for capillary occlusion and hypoxia to be a factor. For the large tumour, we set the cells to be normoxic at the start and, with only five days before treatment, only about 2% of cells were hypoxic and 15% of capillaries occluded by that time. In the next section, we report on treatment results for a tumour of the same size with fully established hypoxia at the time of RL injection.

### Hypoxic resistance

The largely normoxic tumours tested in the previous section illustrate the impact of the escape of radiation from the tumour based on what is effectively a surface area to volume ratio (EDF). We next asked whether established hypoxia and the associated radioresistance could reverse the treatment success observed for the larger tumours in Fig. 3. To test this, we simulated tumours of the same sizes, this time running a pre-simulation procedure, described in the Methods Section, to allow hypoxia to develop before running the full simulation.

The smallest and intermediate tumours were too small to develop hypoxia and so the results for these two were exactly the same as shown in Fig. 3. Over the 40 days of pre-simulation, about 43% of the cells in the large tumour became hypoxic through the process of capillary occlusion limiting the oxygen supply.

Fig. 4 shows the largest tumour being eliminated by the treatment, indicating that hypoxia was insufficient to protect it. Examining the images, we see that the majority of the initial outer layer of normoxic cells and the scattered normoxic cells on the interior turned apoptotic within a few days of the injection. The subsequent images give the appearance of the tumour undergoing an erosive process from the outside inward. The entire tumour was eliminated by Day 26.

**Figure 4.**
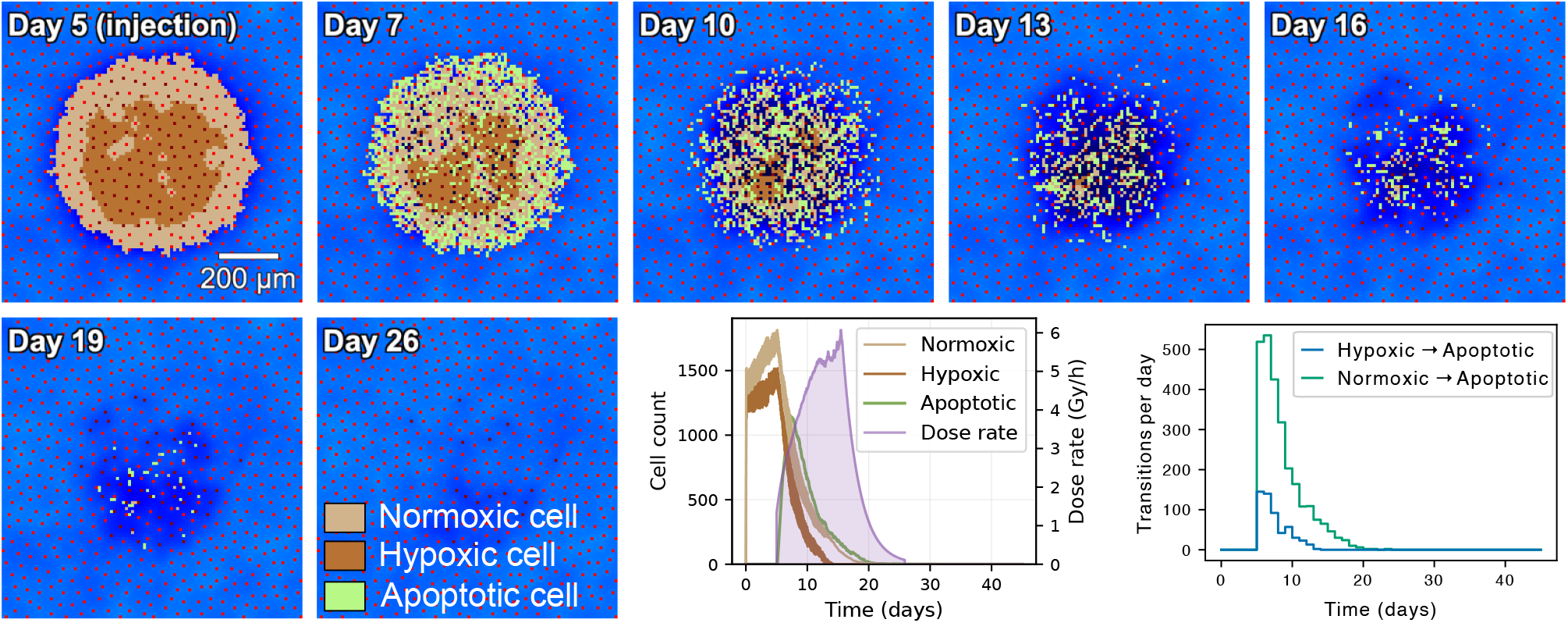
Treatment of a large tumour (initially 600 *µ*m in diameter) with fully developed hypoxic regions. The presence of hypoxic regions did not have the anticipated protective effect on the tumour despite hypoxic cells being more resistant to treatment. However, the time until the tumour was fully eliminated was significantly longer (26 days) than for the normoxic equivalent (16 days). In contrast to the normoxic tumours, which showed rapid uniform apoptosis, here an outer shell of normoxic cells went apoptotic and were subsequently removed, triggering an inward wave of erosion that progressively eliminated the tumour from the outside in. The left plot in the bottom row shows population sizes by tumour cell type, and the dose rate over time. Hypoxic cell counts decrease more rapidly than normoxic ones, disappearing completely by Day 14. The right plot shows the radiation-induced transitions from normoxic and hypoxic to apoptotic. These transition counts show much greater induction of apoptosis in normoxic compared to hypoxic cells, as expected based on hypoxic radioresistance, indicating that the disappearance of hypoxic cells is due to reoxygenation rather than apoptosis induction.

Hypoxic cells were both less radiosensitive and attempted division much less frequently than normoxic cells (at most a 0.05 probability each day) suggesting that this erosion process was driven by reoxygenation of hypoxic cells thereby reverting them to normoxic and increasing their radiosensitivity and division rate. To confirm this reoxygenation explanation, we tracked the number of transitions from normoxic to apoptotic and hypoxic to apoptotic. As expected from hypoxic radioresistance, hypoxic cells became apoptotic at much lower rates (see the bottom right panel of Fig. 4).

We also extended the apoptotic removal time to test whether keeping tumour cells present longer, thereby maintaining capillary occlusion longer, would lead to tumour survival or at least extend its presence. Even with apoptotic cells staying around ten times longer (20 day average removal time), treatment still succeeded and the time to elimination was essentially the same despite the prolonged presence of apoptotic cells. Examination of the simulation output revealed that capillaries did not reopen, as expected, but the drop in oxygen consumption with the change of normoxic cells to apoptotic was sufficient to trigger the inward wave of reoxygenation.

### Quantifying the treatment size threshold and time dependence

The simulations in Fig. 3 suggested that tumour size was a key determinant of treatment success. To quantify this relationship and test its robustness to capillary density (and hence oxygen supply), we swept through a wide range of initial tumour sizes with 150 replicates per size and recorded the average cure rate for each size.

We used an injection amount of 50 nmol to best illustrate the fail/cure transition and for consistency with the two-injection protocol described later. We repeated the sweep at a lower capillary density (420 vs 605 cap/mm^2^) to test the effect of a clinically relevant source of inter-patient variability. Given that the hypoxic tumour in Fig. 4 was eliminated despite hypoxic resistance, we also measured time to elimination to quantify the effect that radioresistance does seem to have. Results are shown in Fig. 5.

**Figure 5.**
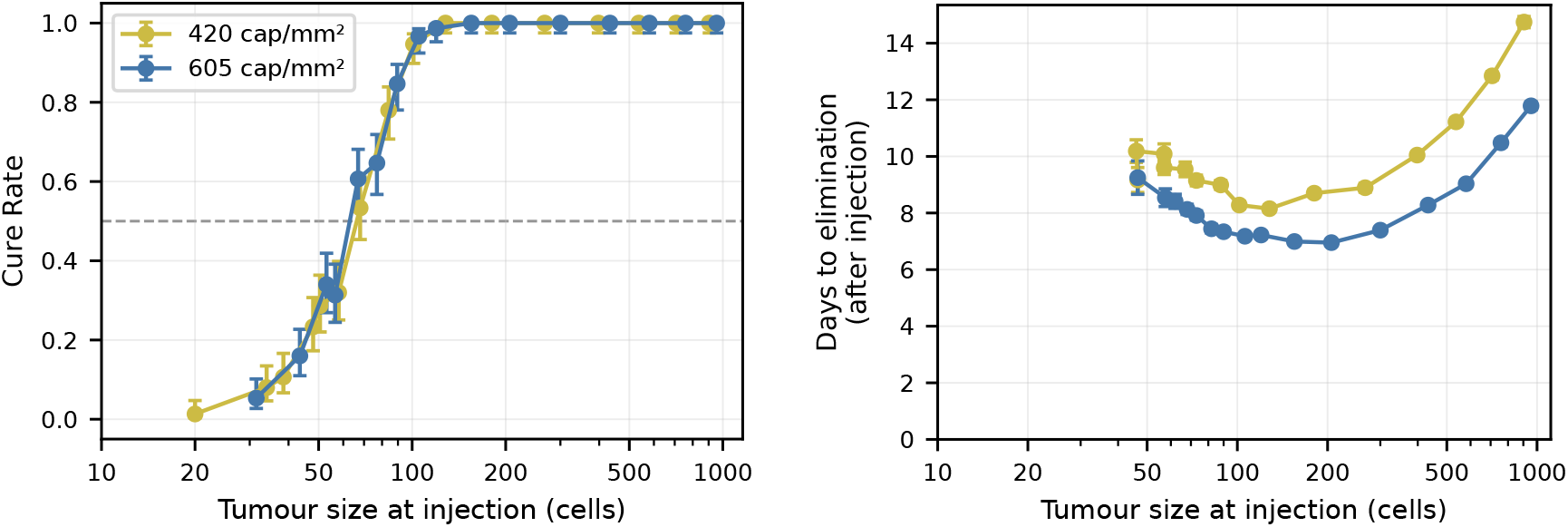
Treatment outcome as a function of tumour size at injection (50 nmol) for two capillary densities (420 and 605 cap/mm^2^). Left panel: Cure rate vs tumour size at injection. The two curves are statistically indistinguishable (with 150 replicates per radius), demonstrating that capillary density has negligible effect on whether treatment succeeds. Error bars show Wilson’s 95% confidence intervals. Right panel: Mean time to tumour elimination (cured runs only) vs tumour size at injection, with error bars showing *±* SEM (with 150 replicates, the error bars are only visible at the smallest tumour sizes). Lower capillary density is associated with consistently longer elimination times consistent with the slower cell cycle (and thus less frequent survival checks) at lower oxygen levels. Data points are omitted where fewer than three cured runs were available, resulting in the elimination-time curves beginning at larger tumour sizes than the cure-rate curves.

The cure-rate curves for the two capillary densities with a 50 nmol injection were nearly identical across the full size range tested (20 to 1400 cell upon injection on day 5), with a transition from failure to success centred around 65 cells and spanning roughly 20 to 100 cells. Treatment success is therefore essentially independent of capillary density and hence of the ambient oxygenation level, at least across the range tested.

We also ran the sweep with a 100 nmol injection (not shown). We found a cure threshold between 14 and 15 initial cells, consistent with the tumour size threshold suggested by Fig. 3 in which a 4-cell tumour survived while a 28 cell tumour was eliminated. Remarkably, the threshold size at injection was essentially the same with the 100 nmol injection as with the 50 nmol injection, a finding explained by the saturation effect explored in the next section. Below approximately 400–450 cells, tumours showed negligible capillary occlusion and negligible hypoxia for both capillary densities, so both curves reflect purely normoxic biology in this range. Above 450 cells, hypoxia and associated radioresistance were present but did not cause treatment failure, consistent with Fig. 4. In the Discussion, we address how incorporating spatially resolved RL delivery — including compromised delivery due to vessel occlusion — might introduce treatment failure at larger sizes into the model [25]. We also address shortcomings of the way DNA damage is tracked in the Lea-Catcheside survival probability that curtail the impact of radioresistance.

The elimination-time curves in Fig. 5 reveal a more subtle story. Both show a clear minimum around 150-200 cells, reflecting two competing effects. At tumour sizes below the minimum, the effective dose decreases due to the EDF and the survival fraction per division attempt rises. Any cell that survives a division attempt, a more likely event at these lower sizes, is replaced by two daughter cells with no accumulated damage, which must themselves sustain damage if the tumour is to be eliminated. This can only occur while RL remains in the system, creating a limited window of opportunity to eliminate these successive generations. Thus a cure takes longer because of additional generations and is possible only within the window of time that the RL is present — a few days to accumulate and a few days of clearance and decay, consistent with the maximum of ~ 10 days seen at the left end of the elimination-time curve. As tumour size decreases further, lower dose rates mean cells have a higher chance of dividing, pushing the opportunities for division attempts beyond the RL window, and the cure rate falls toward zero.

At large tumour sizes, hypoxic cells are present and the erosion process must cycle through successive iterations of reoxygenation, division, and apoptosis from the tumour boundary to the centre. Elimination time therefore increases with tumour radius in this regime, as seen in Fig. 5. The elimination-time curve is higher for lower capillary density most likely because of slowed cell cycle, and therefore less frequent survival checks, at lower oxygen levels.

Both elimination-time curves omit data points where fewer than three cured runs were available, which is why the elimination-time plots begin at larger sizes than the cure-rate plots.

### Treatment success as a function of injected amount and receptor density

Having established that tumour size is an important determinant of treatment success, we asked whether this reflects the size dependence of the EDF, the total receptor content of the tumour, or both, and whether increasing the injected amount could compensate for low receptor density. We simulated treatment of a small tumour (initially 8 cells) over a range of injection amounts and receptor densities. Fig. 6 shows the success rates for each parameter combination by averaging the outcome (failure → 0, success → 1) over 20 replicates. There is a transition zone between treatment failure (dark blue) and success (yellow) identifiable as a noisy gray band that sweeps through the middle of the plot. For injected amounts on the right side of the plot, say above 50 nmol, the colour gradient in the horizontal direction is fairly flat but with a clear transition from low to high cure rate in the vertical direction. This suggests that treatment depends heavily on a patient’s receptor density, at least through the transition, with limited scope for changing the outcome by increasing the injected amount once above 50 nmol.

**Figure 6.**
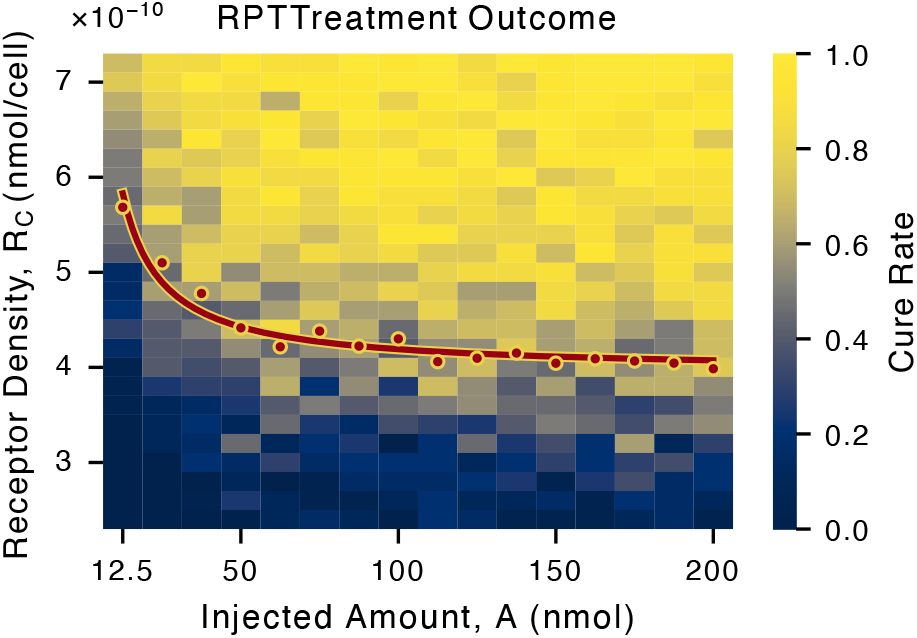
Cure rates for a parameter sweep through injected amounts and receptor densities. Each rectangle represents 20 replicates, simulating treatment of a tumour initially consisting of 8 cells. Treatment success is weakly dependent on total injected amount and strongly dependent on receptor density. The red curve with yellow outline is a fit of Eq. (22) which predicts the locus of a 50% cure rate (red dots). The horizontal axis values in equivalent mCi are 24.5 mCi, 98 mCi, 196 mCi, 294 mCi, and 392 mCi.

Both observations can be understood through analysis of the reduced PK submodel in the Appendix, where we derive the following relationship between the receptor density and the injected amount which can be used to fit the 50% cure-rate level curve in the *A*–*R*_*C*_ parameter plane:

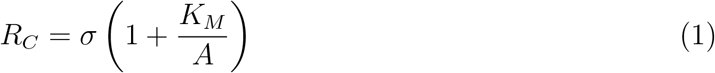

where *σ* and *K*_*M*_ are parameters that we fit and can be related to the parameters of the model.

Without fitting, we can use the parameter values in Table 3 to calculate that *K*_*M*_ = 4.3 nmol. This, as a value that *A* can take on, lies just below the lower end of the injected amount range explored in Fig. 6 (12.5–200 nmol) and below typical clinical injection amounts. This value of *K*_*M*_ suggests that, for values of *A* over 50 nmol (more than 10 times *K*_*M*_), the 50% cure-rate level curve is essentially horizontal (at *R*_*C*_ = *σ*). In fact, the derivation of Eq. (1) applies to any level curve of the cure surface, indicating that above 50 nmol all cure-rate level curves are nearly horizontal — that is, cure rate is essentially independent of the injected amount. For values of *A* below 50 nmol, the cure rates eventually drop to zero as *A* → 0.

For each value of *A*, we fit a logistic function to estimate the 50% cure rate for that slice of the data, yielding the discrete approximation of the 50% cure-rate level curve shown as red dots in Fig. 6. Fitting these points with *σ* and *K*_*M*_ as free parameters (see Appendix) gives *σ* = 3.9 *×* 10^−10^ nmol/cell and *K*_*M*_ = 5.9 nmol (red curve). The fitted *K*_*M*_ is close to the theoretical prediction of 4.3 nmol and well below 50 nmol, supporting the saturation claim. The horizontal asymptote *σ* = 3.9 *×* 10^−10^ nmol/cell marks the boundary below which treatment is unlikely to succeed at any injected amount.

Further interpreting the fit, all simulations above 50 nmol are in the saturating regime where the amount of RL bound and internalized, which we refer to as captive RL and denote by 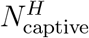, is essentially independent of *A*, explaining the weak observed dependence on injected radioactivity.

The fit also provides a mechanistic picture at the receptor level. At the 50% cure boundary with a typical clinical injected amount *A* = 100 nmol and the fitted value of *K*_*M*_ = 5.9 nmol, *A/*(*A* + *K*_*M*_) = 94.4% of receptors are occupied. Since receptors are nearly fully occupied, additional RL has few available binding sites, whereas increasing *R*_*C*_ directly increases the number of available sites and thus the total captive RL.

Although this fit provides a compelling argument that the saturation of captive RL explains the insensitivity to the injected amount for clinically relevant amounts, the derivation of the fitting curve requires a few rough approximations so we also appeal directly to the pharmacokinetic data from the simulations to confirm the weak A-dependence and strong *R*_*C*_-dependence directly. Fig. 7 shows the time course of 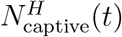 from several of the simulations used to generate the image in Fig. 6. We used three values of receptor density, *R*_*C*_ = 3 *×* 10^−10^, 5 *×* 10^−10^, and 7 *×* 10^−10^ nmol/cell, in blue, green, and red respectively. Each curve shows the average of 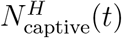 over 20 replicates for a single value of the injected amount. There are 13 curves of each colour corresponding to injected amounts varying from 50 to 200 nmol in increments of 12.5 nmol, each corresponding to a coloured rectangle in one of the three horizontal slices in Fig. 6 at the appropriate *R*_*C*_ value. The peak value for each set of *R*_*C*_ curves is denoted by a dashed horizontal line of the appropriate colour.

**Figure 7.**
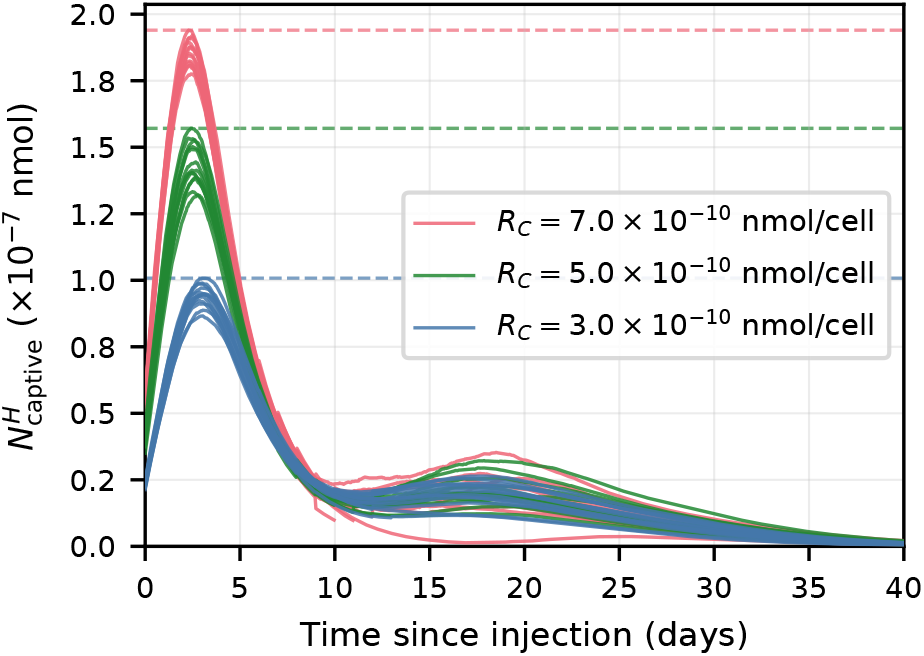
The parameter dependence of cure rate demonstating the impact of RL-receptor saturation. Varying the injected amount from 50 to 200 nmol (curves of the same colour) results in relatively small changes in the amplitude of 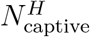 while varying the receptor density (curves of different colors) produces larger changes. This reiterates the observation in Fig. 6 that treatment success is more sensitive to receptor density than to the injected amount.

Consistent with the analytical prediction that, above 50 nmol, we are in a saturating regime with regard to the injected amount, there is little difference among curves of the same colour despite a four-fold change in the injected amount. Also consistent with the analytical prediction is a much larger change in the peaks of the curves between the different *R*_*C*_ values.

We note that the second, lower peak in the curves appears in those simulations for which treatment failed and that have a sufficiently rapid regrowth of the tumour that transiently outpaces the decay and clearance of RL.

### Multi-injection treatment protocols

Single-injection protocols are a useful theoretical baseline, but clinical RPT typically involves multiple injections. We therefore explored how splitting the total injected amount across two injections affects treatment outcome, varying both the time between injections and the relative amounts delivered at each injection.

Fig. 8 shows the success/failure results of treatments as a function of the inter-injection interval, *δ*, and injection skew *S*, with a base injected amount *I*. Two injections were given, separated by *δ* days, where the first amount injected was *I* + *S* and the second amount injected was *I* − *S*. The base amount was *I* = 50 nmol for a total injected amount of 100 nmol as in the previous parameter sweep. We varied the inter-injection interval from 5 to 20 days and the skew from −30 to 30 nmol. We use the word skew here in the sense of shifting part of a distribution from one side of the mean to the other.

**Figure 8.**
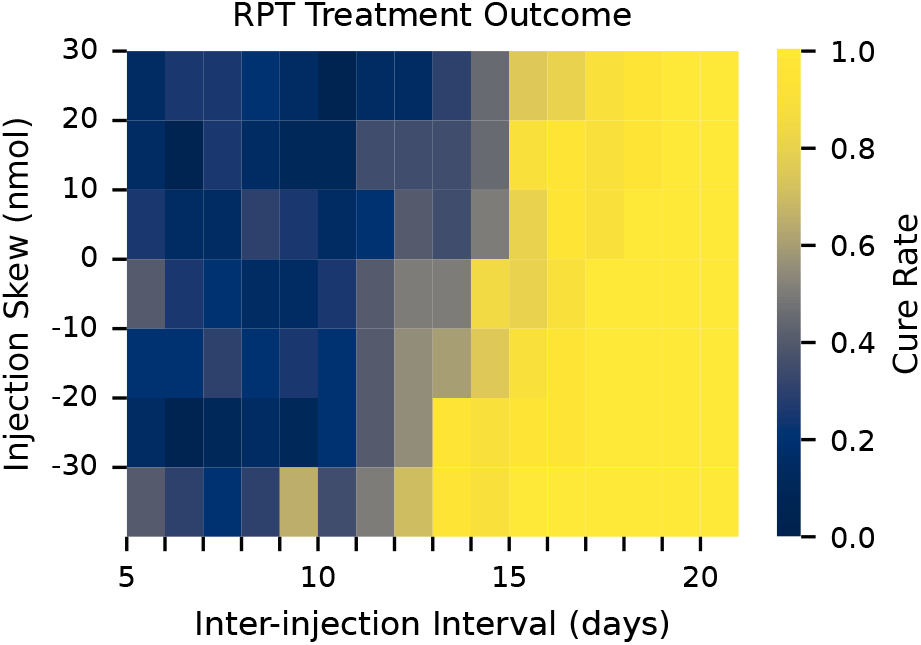
A parameter sweep varying the time between injections in a two-injection protocol and the skew of the injection amplitudes. All treatment simulations included a first injection on Day 5 and a second injection *δ* days later. The first amount injected was 50 nmol + *S* and the second injection was 50 nmol - *S* where *S* is the injection skew. Each rectangle represents the average cure rate over 20 replicates. Tumours initially consisted of a single cell.

As seen in Fig. 8, treatment tends to fail at low inter-injection intervals and succeed at higher intervals with only a slight dependence on the skew. We attribute both of these features to the size-dependence described in Figs. 3 and 5 — if the second injection comes too early, the tumour is still small, the energy deposition fraction is low, and treatment fails. This is reiterated by the fact that positive skew (larger first injection) is associated with slightly lower success rates (the fail-cure boundary tilts a bit to the right). For positive skews, more of the RL is injected while the tumour is smaller and is thus wasted due to the low EDF.

To show that the dependence on inter-injection interval was due to tumour size, we extracted from the interval-skew-sweep data the tumour size at the moment of the last injection (the first injection in cases where that was sufficient to eliminate the tumour and the second injection otherwise) from every simulation with skews from −10 to 10. For each simulation, we noted the tumour size and the success of the treatment. We then binned the tumour sizes, calculated the cure rate for each bin, and plotted the results (see Fig. 9, red curve). We also replotted the curve from Fig. 5 (605 cap/mm^2^) which was a direct assessment of cure rate as a function of tumour size (blue) with an injected amount of 50 nmol.

**Figure 9.**
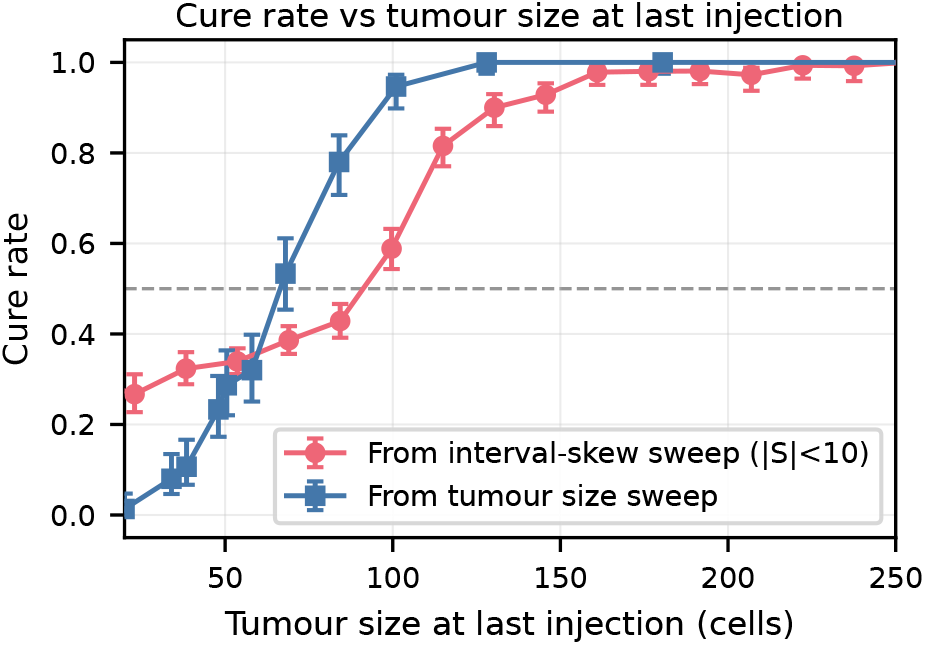
Cure rate as a function of tumour size at the time of last injection. Blue squares: Cure rates are taken from the tumour-size sweep, in which tumours of varying initial size were each treated with a single 50 nmol injection (same curve as in the left panel of Fig. 5). Red circles: Tumour sizes at the time of the last injection were extracted from the interval-skew sweep data (Fig. 8) and binned. The cure rate for each bin was calculated and cure rate was plotted against binned tumour size. Skews with | *S* | ≤ 10 nmol were included (see main text for details). Error bars show Wilson 95% confidence intervals. Both curves show a sharp transition from low to high.

Including different skews in this calculation provided a larger number of “replicates” but introduced the possibility of increasing the variance because of different success rates associated with different injected amounts. However, the low sensitivity to injected amount seen in Fig. 6 suggested this would have minimal impact, a fact borne out by the relatively low variances compared with using only *S* = 0 (not shown).

Both curves show a clear and relatively sharp dependence on tumour size. The baseline cure rate at low tumour sizes is elevated in the interval-skew sweep data. Given that these tumours are small, they must come from instances of small inter-injection intervals and are thus getting an effectively higher injection amount (two of 50 nmol in rapid succession) compared to the size sweep (one of 50 nmol). The failure–cure transitions do not occur at the same sizes (~ 65 cells compared to ~ 100 cells). We attribute this to subtle differences in tumour state arising from the different histories of the two populations, though the precise mechanism remains unclear.

## Discussion

The results presented here can be organized around a single variable: tumour size at the time of injection. Tumour size determines the energy deposition fraction, governing how much of the emitted radiation is absorbed within the tumour rather than lost to surrounding tissue. Tumour size also determines the total receptor content, governing how much radioligand the tumour can capture and internalize. Both effects work in favour of treatment and increase monotonically with tumour size, producing a sharp threshold below which RPT generally fails and above which it generally succeeds. Working against treatment success, larger tumours develop hypoxic cores whose radioresistant cells might be expected to survive treatment and seed regrowth — but in the model this is largely circumvented by reoxygenation, which operates continuously through the protracted action of RPT, even without multiple injections. A more consequential size-dependent obstacle, not captured by the present model, is that the occluded capillaries in larger tumours that cause hypoxia also restrict RL delivery in exactly the regions where radioresistance is highest [30]. Understanding these competing size dependencies — and the conditions under which the beneficial ones dominate — is central to the mechanistic picture developed here.

### General size dependence

The model predicts that, for the specific parameter values explored in our simulations, tumours above the threshold of roughly 65 cells should be eliminated by RPT — a size far below the current detection limit of clinical PET imaging, which is approximately 5–15 mm in diameter depending on lesion contrast and scan time [31, 32]. This suggests that RPT may be particularly effective against micrometastases that are too small to detect or target with other modalities. Because of their narrower DPK, alpha-emitting radiopharmaceuticals have EDFs that drop to zero at even lower tumour radii, suggesting that they would be effective at eliminating even smaller tumours.

### Receptor density as a lever for improving outcomes

As shown in Fig. 6 and explained via the quasi-steady-state analysis in the same section of the Results, treatment outcome depends strongly on receptor density and only weakly on injected amount, at least for amounts in the parameter regime explored in the model. The strong dependence on *R*_*C*_ is made evident in Eq. (20) where it appears as a linear factor in 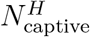. The weak dependence on the injected amount is explained by receptor saturation at *K*_*M*_ ≈ 4.3 nmol. The broader implication is that receptor density is a more productive lever for improving outcomes than escalating the injected amount. Modulating the injected radioactivity is a relatively straightforward option in therapy, and can improve therapy for some tumours of different expression levels [24]. Nonetheless, the observation in this study that modulating receptor content (expression) can have higher impact should be explored further. Although receptor density is not currently a tunable treatment parameter in clinical RPT, patient stratification by receptor expression could be a viable strategy. Tumour-cell specific receptor upregulation, if achievable, could be a worthwhile avenue of research to exploit the sensitivity reported here.

In fact, a related paradigm is already well-known in radioiodine therapy of thyroid cancer, where “redifferentiation” strategies are used to restore expression of the sodium–iodide symporter (NIS) and thereby re-enable radioiodine uptake [33]. In particular, inhibition of the MAPK pathway with BRAF or MEK inhibitors (e.g., Dabrafenib or Trametinib) has been shown to increase iodine avidity in previously refractory disease. Analogous strategies are on the horizon for PSMA RPTs, such as androgen receptor pathway inhibition, and more broadly genetic and epigenetic regulation of receptor expression, offering promising avenues to enhance radioligand uptake and therapeutic efficacy [34]. Future studies should aim to better characterize and leverage receptor upregulation as a means to improve outcomes in radiopharmaceutical therapy.

### Hypoxia and reoxygenation

In the model, RL is not tracked spatially across the extracellular compartment but instead tracked by total amount. Occluded capillaries impact the amount of RL delivered but not in a spatially explicit manner, in contrast with oxygen for which spatial variation in supply does have an explicit impact. Radiation is therefore distributed evenly so that all tumour cells receive the same dose, whether they are hypoxic cells in regions of low oxygen or normoxic cells in regions of higher oxygen. Thus the only mechanism by which hypoxia affects treatment outcome is through reduced radiosensitivity (*α* and *β* values in the Lea-Catcheside survival probability).

In the model, hypoxic radioresistance is overcome by reoxygenation: as normoxic cells on the outer shell of the tumour are rendered apoptotic and eventually removed, local oxygen consumption drops, providing nearby hypoxic cells with an improved oxygen supply, restoring radiosensitivity and accelerating cell division, thereby speeding up induction of apoptosis in these cells. For larger tumours with a substantial hypoxic core, this reoxygenation cycle repeats inward like a wave, consistent with the increasing elimination times seen at large tumour sizes in Fig. 5.

The gradual inward erosion observed in Fig. 4 reflects a combination of three possible processes. First, despite radioresistance, hypoxic cells may still sustain sufficient damage to undergo apoptosis at their next division attempt. Reoxygenation might simply be accelerating the cell cycle on the periphery as oxygen becomes more available with the induction of apoptosis in the outermost (normoxic) cells. Interestingly, this reoxygenation occurs just by induction of apoptosis, and the associated drop in oxygen consumption, and does not even require removal of cells and the subsequent reopening of capillaries, as shown by our lengthened apoptosis-removal simulation. Second, cells sustaining insufficient damage to die as hypoxic cells may reoxygenate, making previously sublethal damage lethal, through the abrupt change in *α* and *β* as cells revert from hypoxic to normoxic — more on this in the next section. Third, reoxygenating cells, now more radiosensitive, continue to accumulate damage from RL that remains in the system. The relative importance of the three processes depends on radiobiological parameters, cell state conversion timescales, and RL clearance rate. Disentangling their contributions would require targeted numerical experiments.

### Model limitations

Two limitations impact the applicability of the model behaviour to real tumours. First, hypoxic resistance is implemented phenomenologically by a change in *α* and *β* when cell types change. In reality, hypoxic cells accumulate less biological damage per unit of physical dose because of reduced oxygen-dependent fixation of DNA damage, and that damage remains unchanged through changes in cell type. In other words, model cells accumulate damage identically but normoxic and hypoxic cells interpret that damage differently upon attempting to divide. Real normoxic and hypoxic cells accumulate damage differently but interpret damage the same way upon attempting division. This means that the second process described above is not relevant in real tumours. This shortcoming would be fixed by a damage model that tracks damage and repair, with different repair rates in different cell types, and uses a single set of survival fraction parameters (*α* and *β*) for both cell types.

Second, in real tumours, occluded capillaries reduce RL delivery to hypoxic regions just as they reduce oxygen delivery, raising the question of whether hypoxic resistance in RPT is primarily a radiobiological or a delivery effect. This makes the model’s size dependence prediction unreliable for larger tumours with occluded vessels. In the model, vessel occlusion begins to appear once tumours reach about 400–450 cells (~ 120*µ*m radius) so our results regarding treatment failure below the size threshold are not influenced by this model limitation. Birindelli et al. [25] demonstrated computationally that in poorly vascularized tumour regions, only a small fraction of radiobiologically hypoxic tissue receives a therapeutic dose of ^177^Lu-PSMA-617, implicating delivery failure as the dominant resistance mechanism in that setting. Since hypoxia and compromised RL delivery are spatially correlated through capillary occlusion, disentangling these experimentally is difficult. A spatially resolved PK submodel, in which RL arrives via open capillaries and diffuses through the interstitium, would address this limitation directly. By separating the radiobiological and delivery effects, the present RL model isolates the radiobiological component and shows that hypoxia-induced radioresistance alone is insufficient to cause treatment failure — a result consistent with, and complementary to, the delivery-focused findings of Birindelli et al.

### Multi-injection treatment protocols

Multi-injection RPT protocols are often motivated by the idea that a second injection timed to coincide with reoxygenation of hypoxic cells could exploit restored radiosensitivity to improve outcomes. In the present model, reoxygenation already plays a role within a single injection — the protracted delivery over several RL half-lives allows successive cycles of hypoxic reoxygenation. In this context, waiting longer between injections is beneficial simply because the tumour grows larger and the energy deposition fraction increases. There is no optimal interval, only a monotone transition from failure at short intervals to success at longer ones (Fig. 8). An optimal inter-injection interval would likely emerge in a model where RL delivery is spatially resolved and sensitive to vessel occlusion, such that a second injection timed too late would find the tumour’s hypoxic core inaccessible to RL. This represents a further motivation for extending the model to include spatial PK. An extended model of this type would also allow us to consider alpha-emitting RPTs which have shorter ranges than beta-emitting RPTs and thus higher EDFs. This makes them more effective against smaller tumors and should make for shorter optimal inter-injection intervals.

### Implications for treatment planning

The results presented here suggest several principles relevant to RPT treatment planning. First, the model predicts that single-injection RPT will fail for tumours below a threshold size at the time of treatment. Rather than waiting for tumours to grow — which risks the formation of new micrometastases in the interim — a more practical strategy would be a two-injection protocol: an initial injection to eliminate all suprathreshold tumours and cut off the source of new micrometastases, followed by a second injection timed to catch previously subthreshold tumours once they have grown above the threshold. The optimal inter-injection interval would be approximately the time required for a single-cell tumour to reach the threshold size. In principle, this could be estimated from tumour growth rate measurements and calibration of the actual size threshold. This provides a mechanistic basis for multi-injection RPT protocols that is distinct from the reoxygenation rationale often cited in traditional radiotherapy.

A second prediction of the model is that receptor density is a more productive target for improving treatment than escalating the injected amount, since receptors appear to be near saturation at clinically relevant injected amounts. Patient stratification by receptor expression — already practised in theranostics — is directly consistent with this finding.

Finally, capillary density affects the speed of treatment response, suggesting that patients with lower vascular density may require a longer observation window before response can be assessed rather than treatment escalation.

Our conclusions drawn from simulations involving only smaller tumours — ones below the threshold where vessel occlusion becomes significant — are robust. Those relating to larger tumours — particularly those in which occluded vessels ought to be restricting RL delivery — should be interpreted with that caveat in mind.

## Methods

### Tumour dynamics

#### Definitions and tumour geometry

The CA consists of a two-dimensional grid of nodes which we treat as a cross section of a cube of tissue 4 mm long on each side. We use the word node to refer to the CA units instead of cells for clarity as not all nodes are biological cells (which we refer to simply as cells). Each node represents a 10 *µ*m by 10 *µ*m object, either a cell or the cross section of a capillary. Each simulation starts with an initial tumour seed of variable size, ranging from one cell to thousands. Cell types include normoxic, hypoxic, necrotic, and apoptotic tumour cells, and healthy cells (represented by empty nodes). Dynamically, necrotic cells do not appear in the simulations, except for the one shown in Fig. 2, due to parameter values and simulation times. Capillaries can be either open or occluded. We explicitly simulate the tumour in 2D but use a conceptual cylindrical extension into 3D for determining (i) the total number of RL receptors in the tumour and (ii) the fraction of the deposited energy that end up within the tumour. This involves counting the number of cells (*T*) currently in the tumour in the 2D simulation domain, determining the radius, *r*_eff_, of a circle with the same area as that many cells by solving 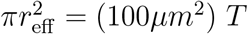, and taking the cylinder height to be 2*r*_eff_. With this geometry in mind, nodes can be thought of as columns of cells of the same type extending into the third dimension and as capillaries penetrating vertically through the tissue block. The circular assumption provides a first approximation to the shape of a tumour in the simulation for those elements that are not modelled explicitly in space; improving upon this would require a detailed spatial accounting of RL in the tumour which we do not do in the present model.

#### Node dynamics

We used the Hybrid Automaton Library (HAL) [35], a specialized Java library designed to manage node dynamics in tumour models. Nodes change between cell types based on the local oxygen partial pressure, with thresholds distinguishing normoxic (high oxygen), hypoxic (mid-range oxygen), and necrotic (low oxygen) states (see Table 1). Transitions are made stochastic by modifying the thresholds by multiplying by a Gaussian with mean 1 and standard deviation 0.04. Radiation can induce a transition from normoxic or hypoxic to apoptotic as described in the radioligand pharmacodynamics section. Apoptotic and necrotic cells do not reproduce, consume oxygen at a negligible rate, and are incapable of converting to any of the other cell type even if oxygen increases locally. Apoptotic cells are removed within a few days and are replaced by healthy cells. Necrotic cells are not removed. See Fig. 1 for a snapshot of node types from a simulation and a legend for the colour encoding used in all figures. The three-tone brown colour bar in the oxygen box to the right of the blue-scale oxygen colour bar indicates the oxygen ranges for normoxic, hypoxic, and necrotic cell types (threshold values in Table 1) with the upper edge of the bar corresponding to the capillary oxygen partial pressure (in Table 2).

**Table 1.**
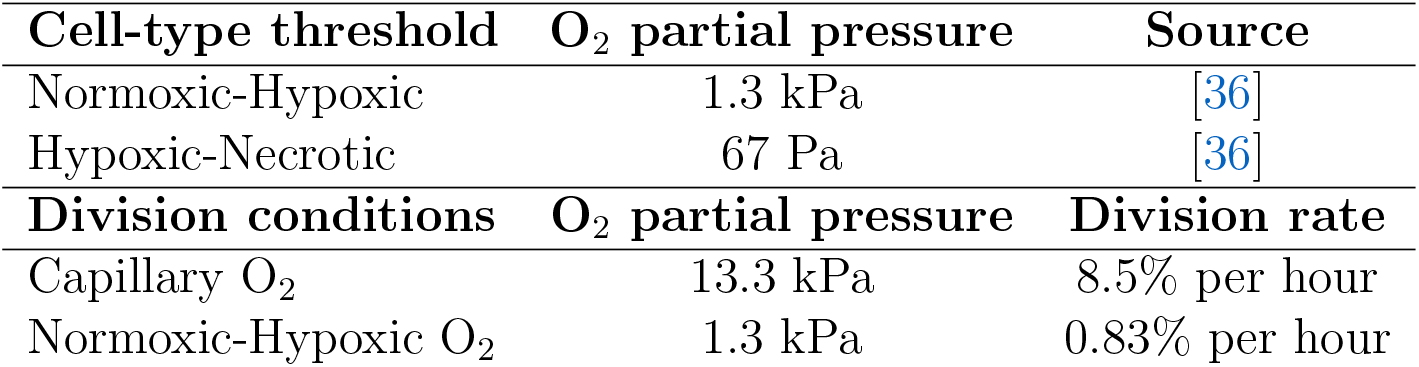
Tumour dynamics parameters. The normoxic-hypoxic and hypoxic-necrotic thresholds were chosen to be consistent with the ranges reported in [36] for radiobiological hypoxia and near-anoxic conditions respectively. Cell division rates decrease linearly as a function of oxygen partial pressure from the maximum rate (at capillary oxygen levels). The maximum division rate was chosen to ensure a high probability of dividing within a day.

**Table 2.** Oxygen flux parameters. The per-area permeability was chosen high enough to get realistic oxygen concentration in regions with only healthy cells but low enough that hypoxia was possible. The apoptotic and necrotic consumption rates were chosen to be negligible compared to the others.

| Oxygen delivery | Symbol | Values | units | Source |
| --- | --- | --- | --- | --- |
| Oxygen diffusion | $D_{O_2}$ | $2 \times 10^{-5}$ | $\text{cm}^2/\text{s}$ | [38, 39] |
| Capillary oxygen pressure |  | 13.3 | kPa | [40] |
| Capillary-tissue transfer rate coefficient | $p$ | 8.3 | $\text{s}^{-1}$ | - |
| Oxygen consumption |  |  |  |  |
| Healthy cell | | 0.4 | $\text{s}^{-1}$ | [41] |
| Normoxic tumour cell | | 2 | $\text{s}^{-1}$ | [42] |
| Hypoxic tumour cell | | 0.8 | $\text{s}^{-1}$ | [43] |
| Apoptotic/necrotic cell | | $3 \times 10^{-4}$ | $\text{s}^{-1}$ | - |

#### Cell division

Each cell type undergoes division at different rates. Normoxic and hypoxic cells can reproduce stochastically into neighbouring “empty” nodes, replacing healthy cells. Division is checked once per hour per cell and each cell is given a probability of dividing based on its type and the local oxygen partial pressure (see Table 1). Because only cells at the tumour boundary have empty neighbours available, growth is surface-driven rather than exponential, and the tumour radius increases approximately linearly in time. This growth model produces an observed growth rate of the tumour radius measured to be around 7 *µ*m/day as can be seen (roughly) in Fig. 2. Division rates vary linearly with local oxygen partial pressure, and since cell type is also determined by oxygen, normoxic cells always divide faster than hypoxic cells. This bias is reinforced by the requirement of an empty node: hypoxic cells, residing deeper within the tumour, rarely have empty neighbours available for division.

As it matters for understanding the reoxygenation phenomenon described in the Results section, we make the order of steps explicit here. A cell first undergoes any changes of state (e.g. normoxic to hypoxic), then there is a stochastic check for division. If division is indicated, there is a survival check as described in the PD section below, and finally a neighbour check to see if a surviving cell that is trying to divide has an empty nearby node into which it can divide.

### Oxygen

#### Vasculature configuration

Capillary positions were generated by placing *n*_cap_ capillaries uniformly at random in the 4 mm *×* 4 mm domain and then iteratively applying a pairwise repulsive force between capillaries closer than a cutoff radius *R*_repel_, with force magnitude proportional to (*R*_repel_ − *d*) (a rectified linear spring), where *d* is the distance between capillaries. Iterations were run for long enough that capillaries were a minimum distance apart but not so long that lattice patterns appeared.

Two CSV files of capillary locations were generated, one for each of two capillary densities: 605 cap/mm^2^ (high density, *n*_cap_ = 9680) and 420 cap/mm^2^ (low density, *n*_cap_ = 6720). The former was chosen to ensure that the capillaries were a few cell lengths apart and that the resulting oxygen distribution supported typical tumour division rates. The latter was chosen to create a lower oxygen background that would not induce hypoxia but was low enough to generate measurable differences in tumour dynamics (e.g. as shown in Fig. 5). A CSV file was loaded at the start of each simulation. The boundary condition for the oxygen PDE was calibrated separately for each configuration as described in the next section.

#### Oxygen influx, diffusion, and consumption

The oxygen partial pressure was tracked in the same 4 mm x 4 mm domain as the CA. We assumed that oxygen enters the domain at capillaries at a rate proportional to the difference between the fixed arterial oxygen pressure *u*_*B*_ and the domain pressure at the capillaries’ node. Consumption occurred at all non-capillary nodes at a rate determined by the cell type. Oxygen in the domain diffused through the interstitial space. The PDE for *u*(*x, y, t*), the oxygen partial pressure at location (*x, y*) at time *t*, is given by

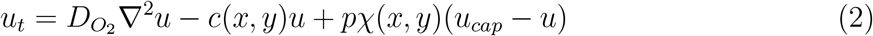

where 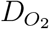 is the diffusion coefficient of oxygen, *c*(*x, y*) is the consumption rate, *p* is the per-area permeability of the capillary walls, *χ*(*x, y*) is the indicator function for open capillaries (zero at cells and occluded capillaries and one at open capillaries) and *u*_*cap*_ is the partial pressure of oxygen in the capillaries. Because the time scale of oxygen diffusion is fast compared to changes in the cell population, we treat the oxygen dynamics as being in quasi-steady state [37]. We set the right hand side of Eq. (2) to zero and solve a discrete version of the resulting PDE using Red-Black Successive Over-Relaxation. The solution to one instance of this PDE is shown in the oxygen panel of Fig. 1 and in Figs. 2, 3, and 4 with the tumour cells overlaid. Parameters used in the oxygen submodel are reported in Table 2.

At the boundaries of the domain, we imposed a non-homogeneous Dirichlet condition, setting the pressure to a fixed value. We determined this value by running a tumour-free version of the model to steady state on the capillary configuration to be used in the actual tumour simulation. We spatially averaged the steady-state oxygen pressure and used the resulting value as the boundary value for the tumour simulation.

We used a simple model to capture the effects of pressure-based occlusion — we counted tumour cells within a 100 *µ*m neighbourhood of the capillary and occluded it if the count exceeded a threshold (50% of the neighbourhood), with a stochastic probability at each time step. Capillaries could reopen in a similar manner but with a lower local-density threshold.

#### Pre-simulation procedure for hypoxia development

To simulate tumours with fully developed hypoxic regions, we used a pre-simulation procedure prior to the main simulation. Starting from an initial fully normoxic tumour seed, we ran the model for 40 days with cell division suppressed and no RL injections. During this period, capillaries surrounded by sufficient tumour cells occluded, and the oxygen field equilibrated to the resulting vascular configuration, allowing hypoxic and necrotic regions to develop. The resulting tumour was then used as the initial state for the full simulation.

### Radioligand pharmacokinetics

The pharmacokinetic component of the full model consists of a compartment model that tracks RL concentration from injection through to internalization within tumour cells. The starting point of the submodel, simplified in the Appendix using time scale analysis, consists of two central vascular compartments (one for hot and one for cold RL) into which RLs are injected, and eight compartments local to the tumour: vascular, extracellular, tumour-cell bound, and intracellular (a hot and cold compartment for each). The tumour vascular compartments, which we refer to as the capillary compartments, are fed by flow from the central vascular compartments and feed into the interstitium, which we refer to as the extracellular compartments (to avoid too many “int” subscripts), by capillary wall permeability. From there, the RLs can bind specifically to tumour cells and can be internalized with their receptors. The amounts in the bound and internalized compartments are together referred to as the amount of *captive RL*. The radioactive isotope portion of internalized RLs can be released from a cell following autophagy of the RL. We treat release as removal from the system instead of adding additional compartments. The compartments track total RL rather than a spatial distribution of RL, an assumption addressed in more detail in the next section. We use the cylindrical extension described above to estimate the total number of receptors in the tumour and use that number in the binding rate. See Fig. 1 for a schematic of this component. Within the PK submodel, there is a wide range of time scales. Using multiple-time-scale analysis, we derived a four-variable reduced system for the slow-changing variables 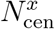, the amount of hot (*x* = *H*) and cold (*x* = *C*) RL in the central vascular compartment, and 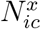, the amount of hot and cold RL in the intracellular compartment. The equations are

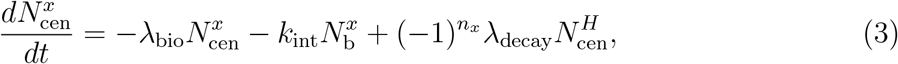

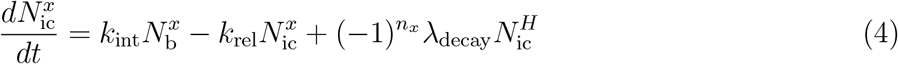

where *n*_*H*_ = 1, *n*_*C*_ = 2 (these exponents on (−1) ensure that radioactive decay moves RL out of the hot and into the cold state), and *N*^*x*^, the amount of RL bound to tumour cells, is

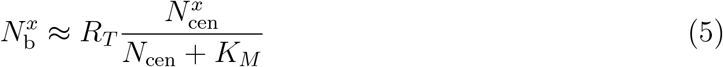

where 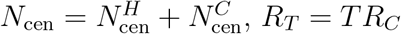, *K*_*M*_ = *V*_cen_(*k*_off_ + *k*_int_)*/k*_on_ = 4.3 nmol, and all other parameters are defined in Table 3. The term *N*_cen_, appearing in the denominator of the Michaelis-Menten-like rate in both ODEs, illustrates the competitive inhibition of cold RL in the binding saturation of hot RL. The other variables are 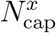, the amount of RL in the capillary compartment, and 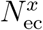, the amount of RL in the extracellular space around the tumour cells. They are in rapid equilibrium with the central vascular compartment:

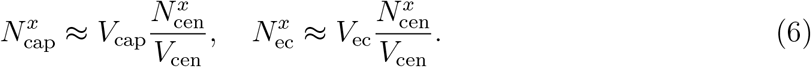

The total number of cells in the tumour, *T*, determines the tumour volume, *V* (each cell is 10^3^ *µ*m^3^), *V*_ec_ is a constant fraction of the tumour volume *V*_ec_ = 0.4*V* and *V*_cap_ is determined by the formula *V*_cap_ = *n*_cap_*A*_cap_*h*_tumour_ where *n*_cap_ is the number of capillaries within the region occupied by the tumour, *A*_cap_ is the capillary cross-sectional area (100 *µ*m^2^) and *h*_tumour_ = 2*r*_eff_.

The multiple-time-scale analysis used to arrive at the quasi-steady state eqs. (3), (4), (5) and (6) is in the Appendix. Parameter values are given in Table 3.

**Table 3.**
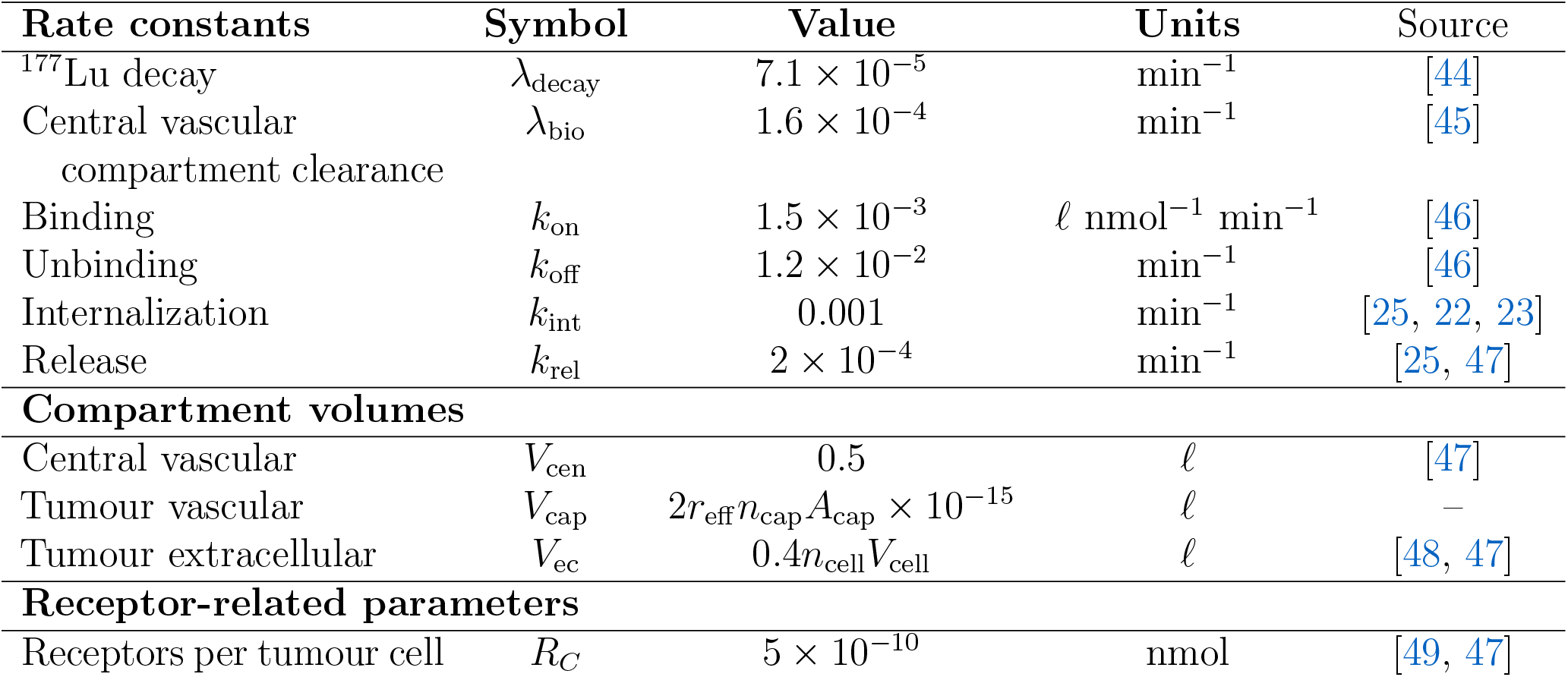
Definitions and values for the parameters in the PK model. The capillary compartment volume is the tumour height (2*r*_eff_) multiplied by the number of capillaries in the tumour (*n*_cap_) multiplied by the cross-sectional area of a capillary. The 10^−15^ converts *µ*m^3^ to litres. The tumour extracellular compartment is 40% of the tumour volume which is the number of tumour cells in the 3D extension *n*_cell_ (all types because this is used to calculate total receptor content) multiplied by the volume of each cell (*V*_cell_ = 1000 *µ*m^3^). *R*_*C*_, the amount of receptors per tumour cell which we refer to as the receptor density, is varied in the receptor-density study but is as given above in all other simulations.

### Injection protocol

Radioligand was injected as a bolus into the central vascular compartment. Each injection consisted of a mixture of hot (radiolabeled, ^177^Lu) and cold (unlabeled) RL in a fixed ratio of 10% hot to 90% cold by molar amount. Both hot and cold RL were capable of binding (competitively) to receptors but only the hot RL delivered radiation.

The injected amount and injection schedule varied by experiment:

- *Single-injection simulations* (Figs. 3 and 4): a single injection of 100 nmol total RL on Day 5.
- *Tumour-size sweep* (Fig. 5): a single injection of 50 nmol on Day 5. This amount was chosen to place the simulations in a regime where the fail-cure transition was clearly visible across the size range tested. It also matches the average injected amounts per injection in the interval-skew sweep.
- *Injected-amount–receptor-density sweep* (Fig. 6): a single injection on Day 5 with total injected amount varying from 12.5 to 200 nmol and receptor density varying from 2.4 *×* 10^−10^ to 7.2 *×* 10^−10^ nmol/cell across the sweep grid. The baseline receptor density used in all other simulations was *R*_*C*_ = 5 *×* 10^−10^ nmol/cell.
- *Interval-skew sweep* (Fig. 8): two injections with a total of 100 nmol. The first injection of 50 + *S* nmol is given on Day 5 and the second injection of 50 − *S* nmol is given *D* days later. The skew *S* ranges from − 30 to 30 nmol and the inter-injection interval *D* ranges from 5 to 20 days.

We have reported the injected amounts in nmol because the RL has a biochemical role in the PK submodel in addition to its radiobiological role, for example, binding to receptors which are measured in nmol. Because we always use 10% hot fraction, the conversion is straightforward — 100 nmol is 7,268 MBq or 196 mCi. This is the most common injected amount used in our studies, with 50 nmol (3634 MBq or 98 mCi) also being used and a range of 12.5 nmol through 200 nmol (908.5 MBq through 14536 MBq or 24.5 mCi through 392 mCi) being the amounts used in the injected-amount–receptor-density sweep.

### Radioligand pharmacodynamics

When ^177^Lu decays, the emitted beta particle can travel up to a few millimeters. The tumours in our model are in the range of 10-600 *µ*m in diameter and the spatial domain is 4 mm across; we do not track the spatial distribution of RLs in the tumour and instead adopt a “well-mixed” assumption — a reasonable approximation given the mm-scale beta-emission range for ultimate dose calculations. We calculate the total amount of hot RL in the tumour and convert that into a dose rate

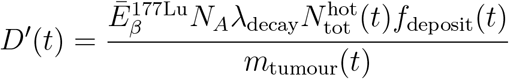

where 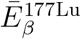 is the average energy of a beta particle, *N*_*A*_ is Avogadro’s number, *λ*_decay_ is the radioactive decay rate, 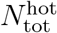 is the total amount of hot RL in the tumour compartments in mol, *f*_deposit_ is the fraction of radiation deposited in the tumour (see the next section), and *m*_tumour_ is the current mass of the tumour.

#### Energy deposition fraction

When RLs concentrated within the tumour decay, the energy is deposited at a distance according to a *dose point kernel* (DPK). By integrating the DPK against a spatial distribution of RL, we can calculate the dose delivered at any point in space in or around the tumour. Some energy will be deposited outside the tumour and this loss of energy is worse for smaller tumours, ultimately going to zero as the tumour radius goes to zero. We refer to the fraction of the total energy deposited that lands within the tumour as *the intratumoural energy deposition fraction* (EDF). In all the simulations presented here, we used a DPK derived from extensive analysis to calculate the fraction (*f*_deposit_) of the energy emitted from RL within the tumour deposited within the tumour [50]. Because we track total RL in each compartment and not the spatial distributions, we use the rough approximation that the tumour is spherical, consistent with the MC-based DPK. Values of *f*_deposit_ are calculated in advance for a set of radii which we use as a lookup table for interpolation on the fly.

Because we relied on unpublished results (the MC simulation), we also used an alternative method to estimate *f*_deposit_ for clarity of explanation and to show qualitative agreement with a simplified approach. In addition to the MC-based DPK, we also used a DPK consisting of a constant value on the interior of a sphere of radius *ℓ* and zero outside the sphere. Because the DPK represents redistribution without loss, the constant value is chosen so that the integral of the DPK over all of space is one. Because the volume of the intersection of two spheres has an explicit formula (found by integrating a surface of revolution), the calculations are exact. We found the value of *ℓ* that gave the best fit of this “uniform-sphere redistribution” DPK model to the MC-based DPK model. The behaviour of the model using the uniform-sphere redistribution DPK model is qualitatively the same as for the MC-based DPK model. Details of the uniform sphere DPK model and the best fit to the MC-based DPK model are given in the Appendix.

#### Survival probability

Throughout a simulation, we keep track of an average state of DNA damage in tumour cells, by age cohort, using a system of ODEs for accumulated dose (*D*), single-strand damage (*A*), and double-strand damage (*G*_alt_). See the Appendix for details. When a cell attempts to divide, it has a survival probability (more specifically a probability of not having unrepaired lethal DNA damage) that is determined by the ODE states:

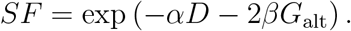

This is a slightly reformulated version (equivalent to the standard formulation) of the Linear-Quadratic (LQ) model with the Lea-Catcheside correction factor [51, 52] to account for ongoing cellular repair. Sensitivity to radiation is modeled by allowing *α* and *β* to depend on cell type, with hypoxic cells more resistant to radiation than normoxic cells. Parameter values are given in Table 4.

**Table 4.** Coefficients for the survival probabilities.

| Coefficient | Value | Units | Source |
| --- | --- | --- | --- |
| $\alpha_{\text{norm}}$ | 0.15 | Gy<sup>-1</sup> | [53, 54] |
| $\beta_{\text{norm}}$ | 0.05 | Gy<sup>-2</sup> | [53, 54] |
| $\alpha_{\text{hypo}}$ | 0.1 | Gy<sup>-1</sup> | [55, 56] |
| $\beta_{\text{hypo}}$ | 0.02 | Gy<sup>-2</sup> | [55, 56] |

**Appendix**

## Pharmacokinetics

Our derivation of the pharmacokinetics component of the model starts with a well-mixed compartment model consisting of ten ODEs for the total amount of hot and cold radioligand (RL) found in each of the compartments: central vascular (cen), tumour vascular or capillary (cap), tumour extracellular (ec), bound to tumour cells (b), and intracellular (ic). Our justification for the well-mixed assumption is based on the fact the typical distance traveled by the emitted beta particles is comparable to or larger than the diameter of the tumours considered. Thus, in a spatial model, any spatial inhomogeneity in the RL distribution would get convolved with a broad kernel to get the distribution of radiation delivery. This smoothing suggests that it is not critical to model the spatial distribution of RL.

The reduced system used in the simulations is described in the main text. Here we present the full system and derive the reduced system from it. The full system is:

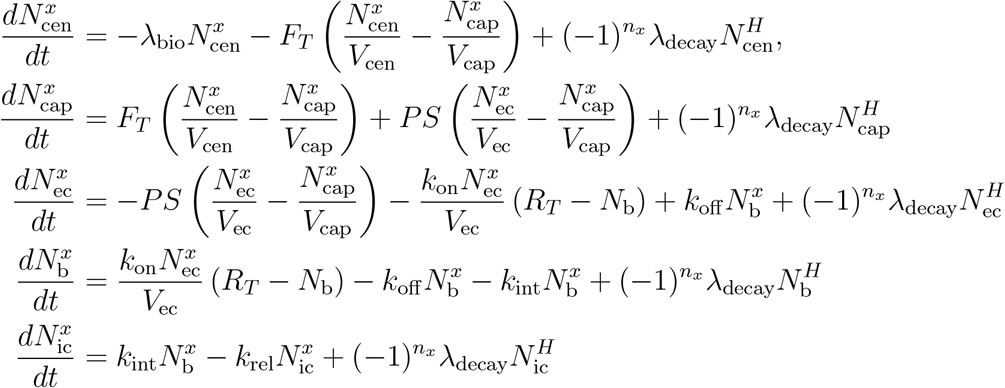

where *x* = *H* for hot/radioactive RL, *x* = *C* for cold/decayed RL, *n*_*H*_ = 1, *n*_*C*_ = 2, and 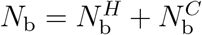 (the total number of bound receptors). The additional parameters not given in Table 3 are given in Table 5.

**Table 5.** Definitions and values for the additional parameters required for the full PK model.

| Parameter | Symbol | Value | Units | Source |
| --- | --- | --- | --- | --- |
| Flow rate between the capillary and central compartments | $F_T$ | $2.9 \times 10^{-3}$ | $\ell \text{ min}^{-1}$ | [47] |
| Permeability rate between capillary and extracellular compartments | $PS$ | $3.5 \times 10^{-3}$ | $\ell \text{ min}^{-1}$ | [47] |

Nondimensionalizing this system, we use 1*/λ*_*bio*_ for the time scale and the typical injected amount of RL, *N*_0_, as the scale for all the dependent variables. Using the substitution *M*_*sub*_ = *N*_*sub*_*/N*_0_ for all the variables, where *sub* is a placeholder for any of the subscripts, we get

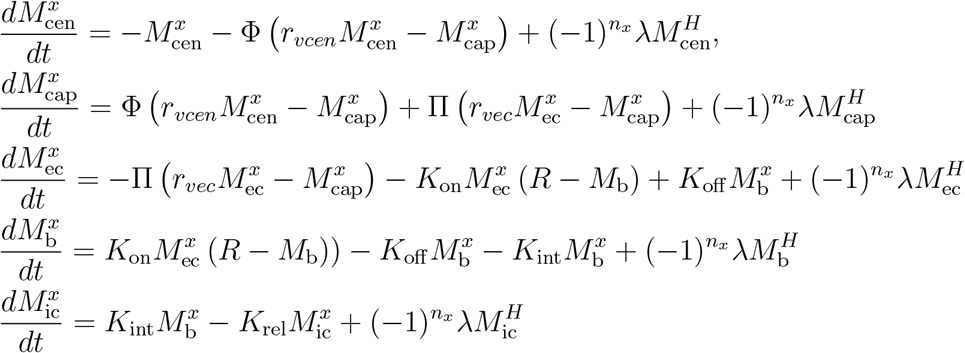

where Φ = *F*_*T*_ */*(*λ*_*bio*_*V*_cap_), *r*_*vcen*_ = *V*_cap_*/V*_*cen*_, *λ* = *λ*_*decay*_*/λ*_*bio*_, Π = *PS/λ*_*bio*_*V*_cap_, *r*_*vec*_ = *V*_cap_*/V*_*ec*_, *K*_on_ = *k*_on_*N*_0_*/*(*λ*_*bio*_*V*_*ec*_), *R* = *R*_*T*_ */N*_0_, *K*_off_ = *k*_off_ */λ*_*bio*_, *K*_int_ = *k*_int_*/λ*_*bio*_, and *K*_rel_ = *k*_rel_*/λ*_*bio*_. Estimating these nondimensional values, we find Φ ≈ 10^8^, *r*_*vcen*_ ≈ 10^−7^, *λ* ≈ 0.44, Π ≈ 10^8^, *r*_*vec*_ ≈ 0.3, *K*_on_ ≈ 10^11^, *R* ≈ 10^−10^ → 10^−4^, *K*_off_ ≈ 10^2^, *K*_int_ ≈ 10, and *K*_rel_ ≈ 1. We can see that the *M*_cap_, *M*_*ec*_, and *M*_*b*_ equations are fast and the *M*_*cen*_ and *M*_*ic*_ equations are slow. The QSS system is

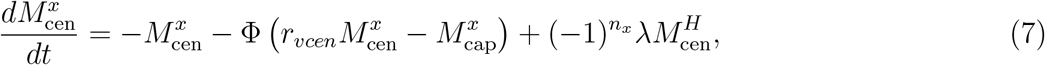

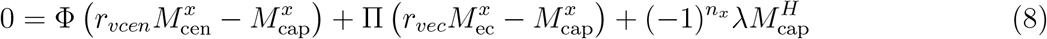

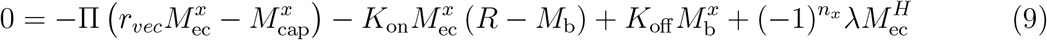

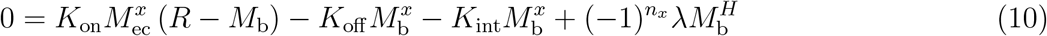

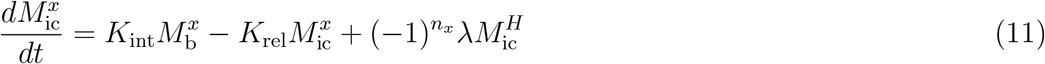

Examining the coefficients reveals that *λ* is at least three orders of magnitude smaller than any other coefficient in the QSS equations so we drop all the radioactive decay terms from equations (8), (9), (10). We find from the nondimensional *M*_cap_ equation, (8), that

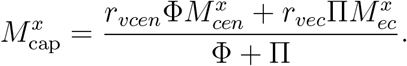

By combining Eqs. (9) and (10), we find that

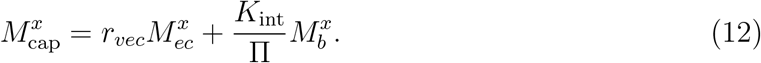

Equating these allows us to eliminate 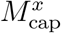, getting

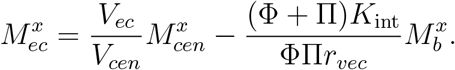

We can rewrite this equation slightly to make better sense of it:

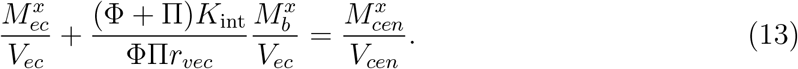

This “concentration equation” says the central concentration is equal to the sum of the extracellular and bound concentrations, with a scale factor in front of the bound concentration. The scale factor in front of 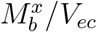 is of order 10^−7^ which suggests that the concentration in the extracellular compartment is essentially the same as in the central vascular compartment.

From both hot and cold versions of Eq. (10), we get a system of equations for 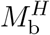 and 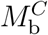 in terms of 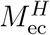 and 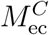:

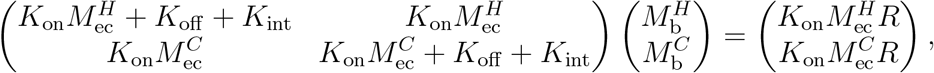

the solution to which is

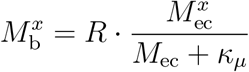

for *x* ∈ *{H, C}* where 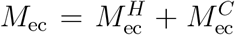 and *κ*_*µ*_ = (*K*_off_ + *K*_int_)*/K*_on_. Plugging this into Eq. (13) gives us

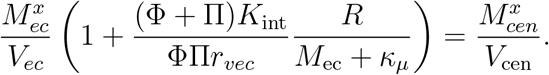

The factor in parentheses is variable because of *M*_*ec*_ but *M*_*ec*_ is at most 1 and *κ*_*µ*_ ≈ 10^−9^ so the entire factor is bounded between 1+10^−12^ and 1+10^−3^. We conclude that 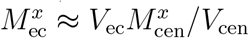 Returning to Eq. (12) and substituting our expression for 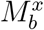, we get

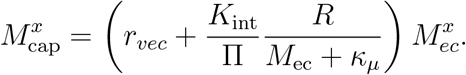

In this expression, *r*_vec_ = 0.3 and the second term in the parentheses is bounded between 10^−13^ and 10^−4^. We conclude that 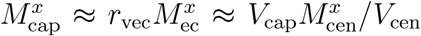. In other words, the tumour vasculature and extracellular space rapidly equilibrate to the same concentration of radioligand as the central vascular compartment. Returning to the 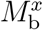 expression above,

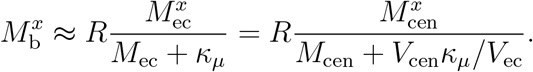

Going back to the ODEs, we can simplify them to

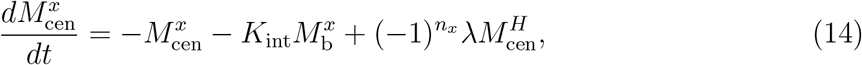

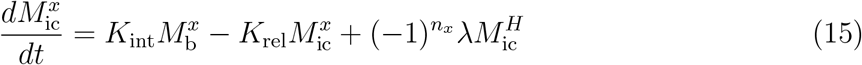

where the 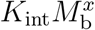 term in the 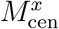 equation comes from chasing the Φ term through the QSS equations.

In dimensional form, the QSS system is

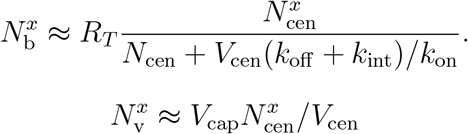

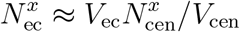

and

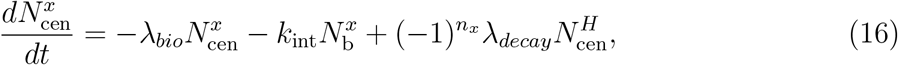

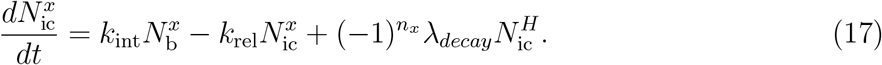

The 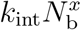 term in the 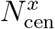 equation is generally much smaller than the 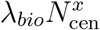 term and could have been left out of this equation but it illustrates the conservation between 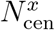 and 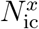. Omitting it allows the *N*_cen_ equations to be solved independently (exponential decay), reducing the system to two equation for 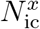.

Notice that 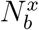 is in the form of number of receptors multiplied by a Michaelis-Menten function with the saturation based on both hot and cold radioligands. This is a mathematical manifestation of the idea that cold ligands act as a competitive ligand to the hot ones.

## Comparison of PK models

The PK submodel initially consisted of a system of 10 ODEs tracking the hot and cold RLs across five compartments. The time scales varied from milliseconds through to days generating significant numerical limitations and motivating the use of multiple time scale methods (quasi-steady state, QSS, analysis). To test that our java implementation of the PK submodel was accurate, we compared it against a separate implementation in python as well as the full 10 variable PK submodel (using a stiff solver) with a tumour of constant size. This confirmed the accuracy of the QSS approximation and its implementation within the full model (see Fig. 10).

**Figure 10.**
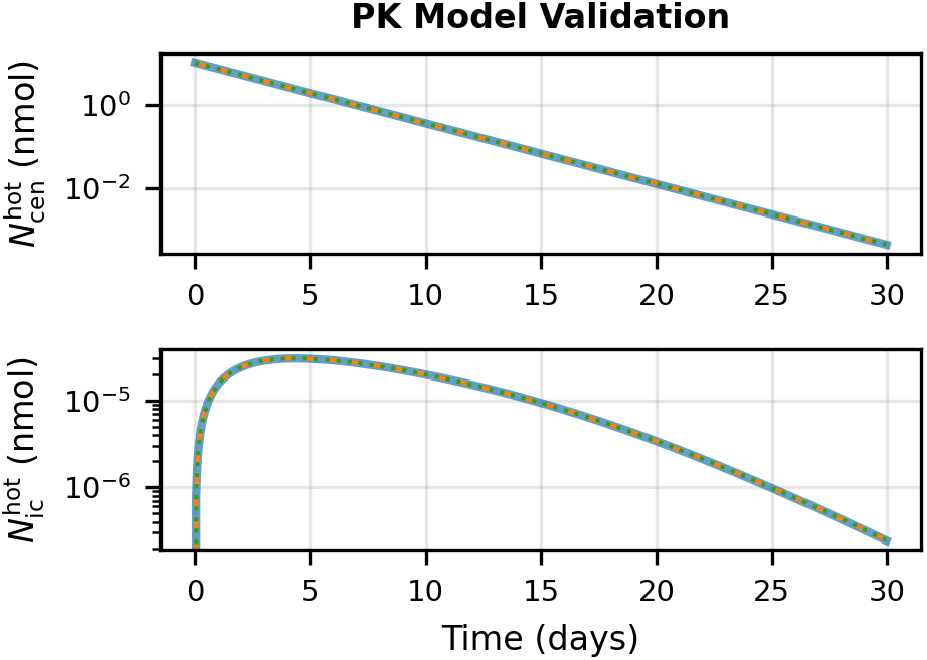
Comparison of three versions of the PK submodel: the full Java simulation with tumour dynamics frozen showing PK dynamics as implemented using the reduced QSS system (blue solid), the full PK submodel integrating all five compartments without the QSS assumption (green dotted), and the reduced PK submodel integrating only the central vascular and intracellular compartments with the others calculated using the QSS (orange dashed). Plots of both central vascular and intracellular compartments. The three solutions are indistinguishable by eye. The full PK system required use of a stiff solver (Python’s BDF implementation of a backward-differentiation solver) for precisely the reason the QSS works (multiple time scales).

## Intratumoural energy deposition fraction

The decision to treat the PK submodel as a well-mixed set of compartments has the benefit of not solving a multi-variable PDE but some useful information is lost. Using a PDE version of the model, finding the deposition of energy from the decaying RL throughout the tumour would require convolving the RL distribution with a kernel that encodes the average redistribution of energy from emission to deposition. For beta particles, this kernel would be a function that decays with distances and has an average redistribution distance of about 0.25 mm. Without the spatial information, the crudest approximation would be to assume that all the energy emitted gets deposited uniformly throughout the tumour. This could miss a lot of detail, especially for a tumour with a complex geometry. We implement a simple correction to recover at least one feature of the ideal calculation. The uniform assumption does not account for the fact that some of the emitted energy will land outside the tumour. For a small tumour, this can be dramatic. An extreme example is a tumour consisting of a single cell — a huge portion of the emitted energy would get spread to the surrounding healthy tissue. Fig. 11 illustrates how energy loss scales with the size of the tumour.

**Figure 11.**
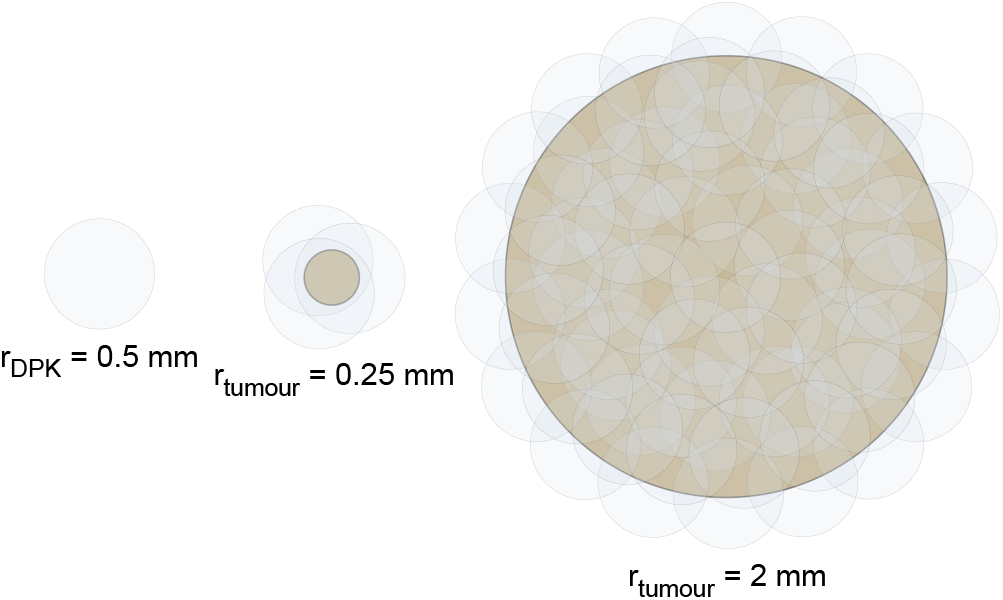
Illustration of the dependence on radius of the Energy Deposition Fraction (EDF). Two tumours are shown (light brown), one with radius 0.25 mm and the other with radius 2 mm. The Dose Point Kernels (DPK) for the RLs are shown as nearly-transparent outlined circles, representing our simplified DPK with a uniform deposition of beta particles across a circle of radius 0.5 mm. The larger tumour has more RL distributed across a larger area, with RL proportional to the area, suggesting that cells within both tumours would receive the same dose. In fact, with the finite cutoff in this DPK, all cells inside the tumour more than 0.5 mm away from its edge do receive the same dose. However, close to edge, some particles leave the tumour. For the smaller tumour, all points within the tumour are within 0.5 mm of the edge, thus receiving a smaller dose than most of the cells in the larger tumour. The fraction of the beta particles or equivalently of the deposited energy that remains in the tumour is the EDF.

To account for this loss of energy, we used results obtained by collective analysis and fitting [50] of multiple calculations of beta particle decay from tumours of varying sizes. We subsequently obtained a dose point kernel (DPK) which characterizes the redistribution of energy from emission to deposition and from that an intratumoural energy deposition fraction (EDF). In the Supplemental Information, we illustrate the idea using an abstract formulation that assumes the DPK takes a simple form that allows exact calculations.

## ODE formulation of the Lea-Catcheside G-factor

The Lea-Catcheside factor *G* accounts for repair of sub-lethal damage during protracted radiation exposure. Traditionally expressed in terms of nested integrals, it can be equivalently formulated as a system of ordinary differential equations (ODEs). This reformulation is mathematically identical but computationally more efficient, conceptually clearer, and more straightforward to generalize.

The survival fraction for protracted radiation delivery, as expressed by Lea and Catcheside, is

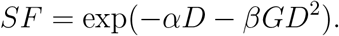

The factor *G* is

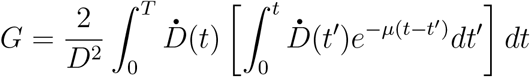

where *µ* is the repair rate constant, *T* is the total irradiation time (h), 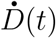 is the dose rate at time t in Gy/h, and 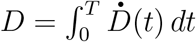 *dt* is the total dose in Gy.

The *SF* can also be expressed as

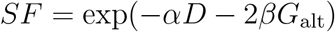

where the factor *G*_alt_ is:

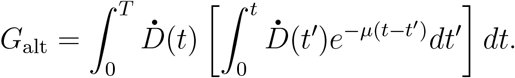

While dividing and multiplying by *D*^2^ makes the connection to the LQ model more explicit, this is a somewhat artificial means of illustrating the connection. Examining the integrals reveals the hidden squared dependence on the dose rate so the relationship is genuine but more subtle than a simple *D*^2^ factor. Expressing it in the latter format avoids both artifice and an unnecessary division by *D*^2^ which is impractical for the purpose of numerics when *D* is small or zero.

Simulating this by quadrature is *O*(*n*^2^) so we reformulate it as an ODE, interpreting the integral terms as quantification of DNA damage. First, we define a function that tracks the amount of single-strand breaks in the DNA of a cell:

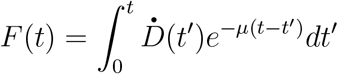

and take its derivative:

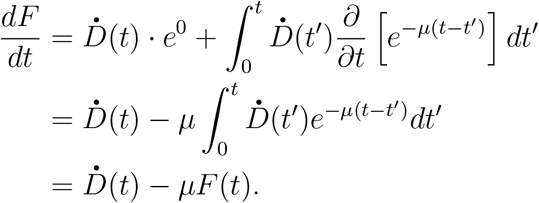

In words, single-strand breaks accumulate at a rate given by the incoming radiation 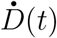 and are repaired with rate constant *µ*.

We define a second function that tracks the amount of double-strand breaks in a cell:

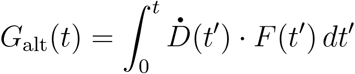

which is the integral factor in *G*. For the ODE formulation, we take its derivative

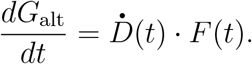

With this ODE formulation, we only need to keep track of the current state of DNA damage and exposure (*D, F, G*_alt_). We do this for each age-cohort (all cells born on the same day) since each age cohort shares an exposure history and interpret these as population averages. When any particular cell attempts to divide, to determine if it survives or goes apoptotic from DNA damage, we use *G*_alt_ and *D* to calculate *SF*, as above, and check this probability against a random number (*U*([0, 1])).

Although we do not implement it in our current model, this ODE formulation opens the option of specifying different repair rates for different cell types (normoxic, hypoxic) or for different phases of the cell cycle, eliminating the need for different values of *α* and *β* in *SF* for normoxic and hypoxic cells.

## Quasi-steady-state analysis of captive RL

We define captive RL as 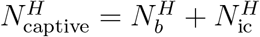, the hot-fraction RL directly associated with the tumour through binding or internalization. Starting from the QSS expression for the bound compartment (Eq. (5)),

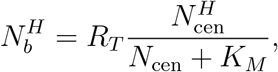

and assuming 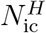 is at quasi-steady state relative to *N*_cen_, a simplifying assumption that is not strictly satisfied but provides a useful closed-form approximation,

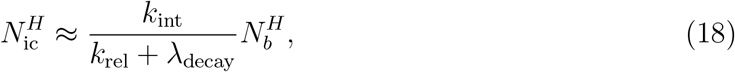

we obtain

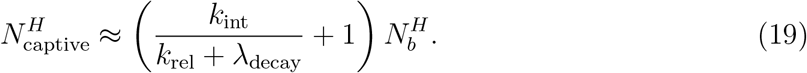

Omitting the − *k*_int_*N*_*b*_ from Eq. (16) (because *N*_*b*_ ≪ *N*_cen_ during the peak period), we get that the central vascular compartment decays approximately exponentially:

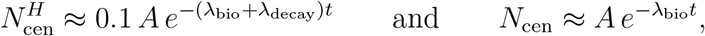

where *A* is the initial injected amount. Substituting these into Eq. (5) gives

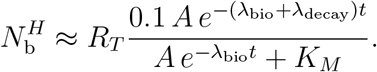

Using this to expand Eq. (19), we get

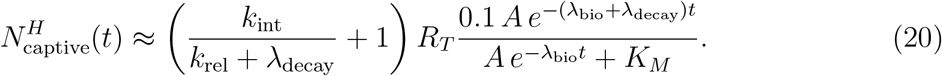

This approximation to 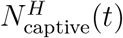 is monotone decreasing in time (for a fixed tumour size) and so evaluating it at *t* = 0 gives a reasonable value for gauging the effectiveness of treatment. At *t* = 0 we get that the captive RL amount is a Hill function in *A* with half-maximum at *A* = *K*_*M*_:

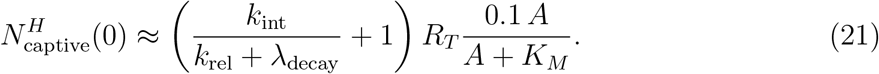

We can replace *R*_*T*_ by *T*_0_*R*_*C*_, where *T*_0_ is the tumour size upon injection and *R*_*C*_ is the amount of receptors per cell. Defining 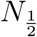 as the (unknown) captive amount that ensures a 50% cure rate, we can set 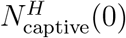 equal to 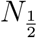 (or use any other subscript between 0 and 1 to get other level curves) and derive an expression for the *R*_*C*_–*A* curve along which the cure rate is 50%:

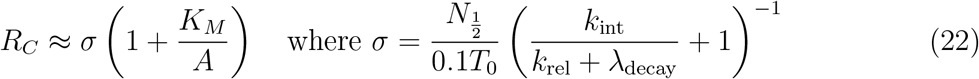

where *A* is the injected amount and *K*_*M*_ = *V*_cen_(*k*_off_ + *k*_int_)*/k*_on_. We fit this expression to the half-cure points calculated from the *R*_*C*_–*A* parameter sweep (see Fig. 6 for the data and the next section for the methods) to get values for *σ* and *K*_*M*_ and then calculate 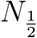 using *σ* and the other parameters in the *σ* expression above.

There are two sources of error in this approximation. First, the 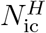 quasi-steady-state assumption ignores the fact that *N*_cen_ and *N*_ic_ change on the same time scale. Second, *R*_*T*_ is calculated as the product of *R*_*C*_ and the number of cells in the tumour. Although we have assumed nothing about *R*_*T*_ in this calculation, the fact that it changes in time, possibly growing, possibly shrinking, depending on the response to treatment, the saturation argument strictly applies only at *t* = 0 and becomes messier with changes in tumour size. As explained in the main text, appealing to the simulation data, using the argument here as a guide on what variables to examine, we confirm that this prediction is at least qualitatively correct.

## Fitting the 50% cure boundary to the injected-amount–receptor-density sweep

Eq. (22) defines a one-parameter family of curves in the (*A, R*_*C*_) plane, parameterized by *σ* and *K*_*M*_, along which the predicted cure rate is constant. To fit this curve to the 50% cure-rate level curve of the simulation data in Fig. 6, we extracted the empirical 50% cure boundary from each column of the heatmap and then performed a least-squares fit to the full collection of points.

For each value of injected amount *A*_*i*_, the column of the heatmap gives cure rates at discrete receptor densities. We fit a logistic function

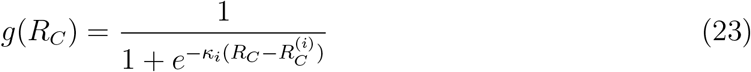

to these cure rates by nonlinear least squares, where 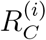 is the receptor density at which the fitted logistic equals 0.5 and *κ*_*i*_ *>* 0 is the steepness parameter for column *i*.

This yields a set of points 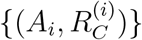 representing the model’s empirical 50% level curve. We then fit *σ* and *K*_*M*_ using python’s scipy.optimize curve fit function, yielding *σ* = 3.9 *×* 10^−10^ nmol/cell and *K*_*M*_ = 5.9 nmol.

We also tried fixing *K*_*M*_ at the theoretically predicted *K*_*M*_ = 4.3 nmol and fitting *σ* by minimizing the sum of squared residuals, which has the closed-form solution

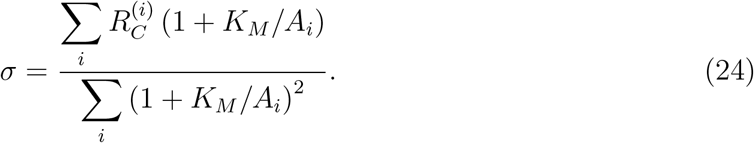

This yielded *σ* = 4.1 *×* 10^−10^ nmol/cell, fairly close to the two-parameter fitting result.

With *σ* = 3.9 *×* 10^−10^ nmol/cell, *T*_0_ = 8 cells, and (*k*_int_*/*(*k*_rel_ + *λ*_decay_) + 1) = 4.69, we get

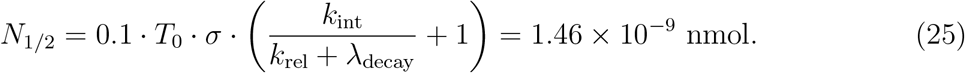

## Acknowledgements

We acknowledge funding support by the IAEA Marie Sklodowska-Curie Fellowship Programme (MSCFP) scholarship, the BC Cancer Foundation, and the Canadian Institutes of Health Research (CIHR) Project Grant PJT-162216. We also acknowledge helpful discussions with Dr. TJ McColl.

## Data and Code Availability Statement

The code for the model is publicly available on GitHub: https://github.com/ecytryn/TumourRPTmodel

## Supplemental Information

### A simple model of the energy deposition fraction (EDF)

We model the fraction of the energy from Lu-177 beta particles emitted from within the tumour that is deposited back onto the tumour itself using a simplified geometric approach. Our model makes the simplifying assumption that each radioactive decay in the tumour emits a beta particle that, on average, deposits its energy uniformly within a sphere of radius *ℓ* (roughly twice the mean distance travelled) around the emission point, and that the tumour is spherical with radius *R* and contains uniformly distributed RL. Although for other reasons, our 3D extension of the tumour was cylindrical, its height was 2*r*_eff_ so a sphere captures the same basic scaling.

Consider a tumour containing *N*_hot_ atoms of Lu-177 uniformly distributed throughout its volume, 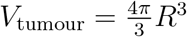. The activity in a small volume 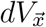 at position 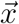 within the tumour is

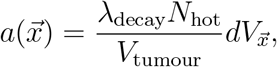

where *λ*_decay_ is the Lu-177 decay constant. Each decay from this small volume deposits energy uniformly over a sphere of radius *ℓ* centered at 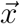, with volume 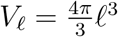. The rate at which particle energy is deposited in a small target volume 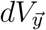 at position 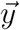 from the small volume 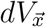 at position 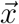 is therefore

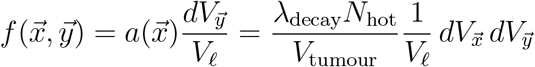

when 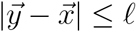, and zero otherwise. The rate of particle deposition in volume 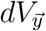 from the activity everywhere in the tumour is

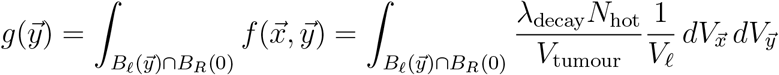

where 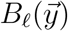 is a sphere of radius *ℓ* centered at 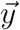 (all points 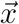 within a distance *ℓ* of 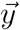) and *B*_*R*_(0) is a sphere of radius *R* centered at the origin (all points inside the tumour). The integrand is constant on the intersection so the integral is the integrand multiplied by the volume of the intersection and so the rate of particle deposition in a small volume at point 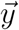 from all sources within range can be expressed as

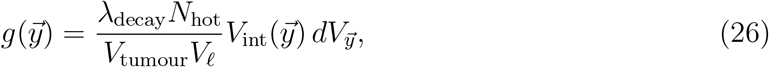

where 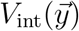 is the volume of the intersection 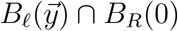.

To obtain the total energy deposition rate within the entire tumour, we integrate over all points 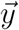 inside the tumour, giving

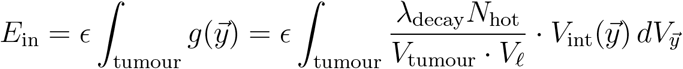

where *ϵ* is the mean beta particle energy per decay. The total energy emission rate from all decays in the tumour is *E*_total_ = *λ*_decay_*N*_hot_*ϵ*. The intratumoural energy deposition fraction, defined as the ratio of the total energy deposition rate in the tumour to the total energy emission rate, is

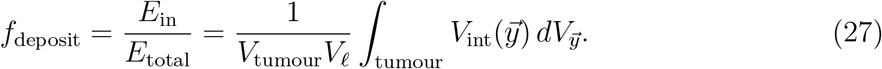

Exploiting the spherical symmetry of the problem, we note that 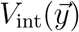 depends only on 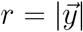, the distance from the tumour center. Converting to spherical coordinates, we obtain

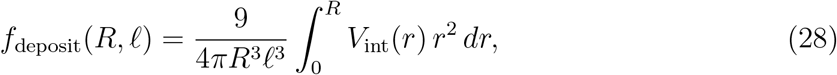

where the factor out front comes from the two volumes in the denominator and a 4*π* in the numerator from the spherical integration. When the spheres do not overlap (*r* ≥ *R* + *ℓ*), we have *V*_int_(*r*) = 0. When one sphere is completely contained within the other (*r* ≤ | *R* − *ℓ* |), the intersection volume equals the volume of the smaller sphere, 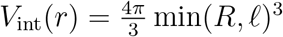<monospace>. For the case of partial overlap 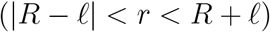, the intersection volume is given by a standard formula for sphere-sphere intersection:

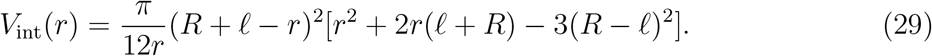

For sufficiently small tumours (*R < ℓ/*2), the tumour sphere is always completely contained within the deposition sphere of radius *ℓ* centered at any point within the tumour. In this regime, *V*_int_(*r*) = 4*πR*^3^*/*3 for all *r* ∈ [0, *R*], and the integral simplifies to *f*_deposit_(*R, ℓ*) = *R*^3^*/ℓ*^3^. Thus, not only does the EDF go to zero with the size of the tumour, it does so like *R*^3^.

Using numerical integration to calculate the EDF, *f*_deposit_(*R, ℓ*), for tumours of radius *R* between 0 and 3 mm, we fit the EDF derived from the MC-based DPK with the EDF derived from the Uniform Sphere DPK (27) with a single optimization parameter (*ℓ*) and found an optimal value of *ℓ*^*^ = 0.41 mm.

Fig. 12 shows *f*_deposit_(*R, ℓ*^*^) (solid blue curve) and the EDF derived from the MC DPK model (red dots) to which it was fit. For reasons that are as yet unclear, it appears as if the EDF derived from the MC-based DPK does not approach zero as rapidly as our more abstract EDF. In the RPT simulations, we use the MC-based DPK model results by linearly interpolating values from a lookup table consisting of the coordinates of the red dots. For the extrapolation from *R* = 0.1 mm down to 0 mm, inspired by the results above, we used a cubic of the form *cR*^3^ where *c* was chosen to go through the value at *R* = 0.1 mm.

**Figure 12.**
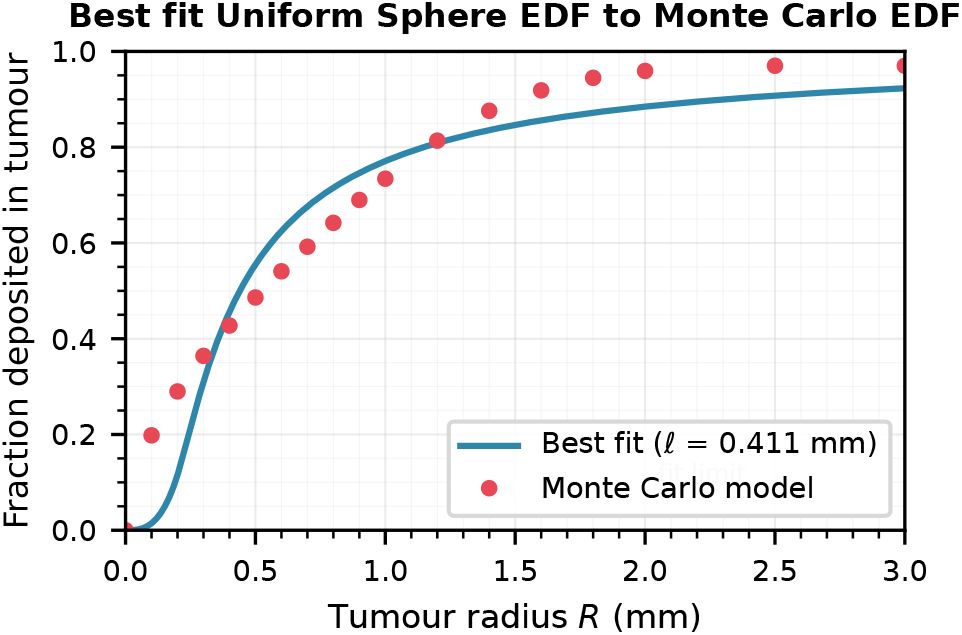
The EDF derived from the MC DPK (red dots) and the best fitting EDF, *f*_deposit_(*R, ℓ*^*^), derived from the Uniform Sphere DPK (blue) with optimal radius *ℓ*^*^ = 0.41 mm. The fit is generally within an order of magnitude except near the origin (*<* 0.3 mm). This deviation suggests that using the best-fit uniform sphere model for the full simulation would be qualitatively similar but would differ quantitatively in the details, including the threshold tumour size at which treatment switches from failure to success.

